# Selective suppression and biasing of chemokine receptors CCR9 and ACKR4 through targeting CCL25 with de novo miniproteins

**DOI:** 10.64898/2026.01.23.701245

**Authors:** Bas de Boer, Thomas D. Lamme, Karlijn Verdwaald, Sara Santamaria Medina, Csongor G. Németh, Elisabeth M. Elfrink, Martine J. Smit, Iwan J.P. de Esch, Christopher T. Schafer

## Abstract

Chemokines and their receptors mediate cell migration and coordinate immune responses, while dysregulation can lead to inflammation. Therapeutic modulation of the chemokine signaling axis has proven difficult. Most drug discovery efforts target the receptors, whereas natural regulatory mechanisms modulate immune responses by directly binding to or modifying the chemokine ligands. Despite this insight, development of effective chemokine-directed modulators has remained elusive. Recent advances in de novo protein design offer an unprecedented opportunity to produce high-affinity binders that efficiently block protein-protein interactions. We implemented a computational workflow leveraging the BindCraft platform to generate miniprotein binders against CCL25, the chemokine ligand for the receptors CCR9 and ACKR4 and implicated in inflammatory bowel diseases. The unbiased development results in several miniproteins designed to block the receptor N-terminus from wrapping the chemokine and prevent productive engagement. Thus, these proteins suppress CCL25-mediated effector coupling and halt MOLT-4 lymphoblast migration. Another class of miniprotein, represented by VUP25111, is predicted to bind CCL25 along the chemokine β1 strand and retained receptor binding. This complex inhibited arrestin recruitment to CCR9, but not to ACKR4, indicating receptor specificity. Additionally, G protein recruitment to CCR9 was unimpeded by VUP25111, suggesting that the miniprotein biased the native agonist towards G proteins. These results demonstrate the effectiveness of differentially targeting CCL25 to suppress CCR9 signaling and new tools to resolve the structural basis of chemokine receptor activation and bias.

## Introduction

Chemokines are small secreted proteins (∼10 kDa) that direct cell migration throughout the body by activating chemokine receptors, a subfamily of G protein-coupled receptors (GPCRs). The chemokine system plays critical roles in coordinating immune responses and homeostasis, as well as directing cell positioning during development and wound repair [1]. Dysregulation of either the receptors or chemokines results in inflammatory diseases, cancer metastasis, and immune evasion of cancers and pathogens [2]. While attractive therapeutic GPCR targets, it has proven difficult to develop effective drugs towards chemokine receptors and only a few candidates have reached the clinic. This suggests that new targeting strategies are needed to fully exploit this versatile and promising signaling pathway [3,4].

The receptor:chemokine system comprises ∼50 chemokines and ∼20 receptors, forming an intricate web of cross-reactivity that allows for highly specific cell positioning [5]. Chemokine binding to chemokine receptors is often simplified to a two-site, two-step model, predominantly involving contacts with the receptor N-terminus and the orthosteric ligand-binding site in the transmembrane (TM) domain, termed chemokine recognition site (CRS) 1 and 2, respectively (**Fig. 1A**) [6–8]. Early models proposed that CRS1 would catch the chemokines and orient binding into CRS2 for receptor activation. Subsequent studies suggested that the ligands interact first with the TM domain of CRS2, and the N-terminus follows by wrapping around the chemokine to stabilize a higher affinity state [9]. Both sites are generally required for activation, with N-terminal receptor truncations or CRS2 point mutations being equally effective at impairing signaling responses [9–12]. Proper binding leads to GPCR activation and coupling of G protein and/or arrestin effector proteins.

**Figure 1.**
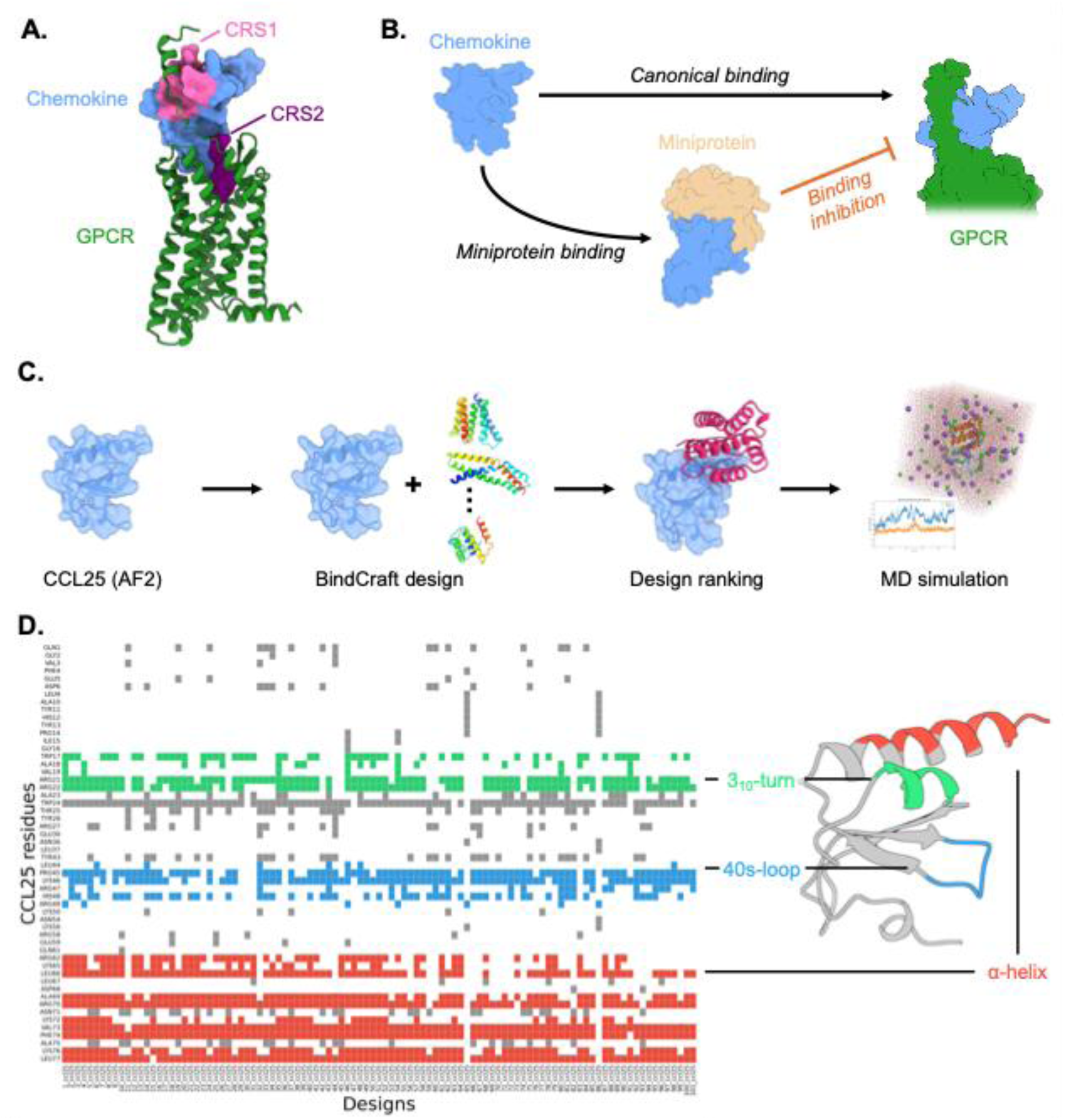
Design strategy and computational workflow for miniprotein binder design against CCL25. (**A**) Chemokine interacts with chemokine receptors primarily through CRS1 (Pink, receptor N-terminus with chemokine body) and CRS2 (Purple, chemokine N-terminus with receptor TM domain). The receptor:chemokine complex is generated using AF3 with sequences of CCR9 and CCL25. (**B**) Conceptual framework illustrating the strategy for inhibiting canonical chemokine to receptor binding by directly targeting chemokine with designer miniprotein binders. (**C**) Computational workflow used to generate chemokine-binding miniproteins. (**D**) Molecular interaction map summarizing contacts between all miniprotein designs and CCL25. The most frequently targeted structural elements, the α-helix, the 3_10_-turn, and the 40s-loop, are highlighted in red, green, and blue, respectively.

The CC chemokine receptor 9 (CCR9) is expressed on thymocytes and intestinal lymphocytes [13,14] and promotes the migration of immune cells between the thymus and small intestine in response to its sole native agonist, CC chemokine CCL25 [15,16]. This receptor:chemokine pair regulates T cell development in the thymus and coordinates immune homeostasis and surveillance in the colon. Thus, overexpression and imbalance of this signaling axis are implicated in inflammatory bowel (IBD) and Crohn’s diseases [17,18], as well as contributing to cancer metastasis to the gut [19,20]. Clinical development of CCR9 antagonists has thus far been unsuccessful, with efficacy in patients remaining a significant challenge [21].

Biological interference of the receptor:chemokine axis preferentially occurs at the level of the chemokine rather than the receptors. For example, pathogens and pests modulate immune responses targeting chemokines. To avoid the host’s immune response, ticks have evolved small, salivary chemokine-binding proteins called evasins that block the chemokine interactions with receptors or glycosaminoglycans [22,23]. This action prevents the immune system from responding and impeding the tick’s blood meal. Another, perhaps more cumbersome, way to target chemokines is by atypical chemokine receptors (ACKRs) that scavenge chemokines by internalizing the receptor:chemokine complex into the cell, thereby altering the chemokine gradients that are essential for chemokine-driven cell migration [24]. For example, CCL25 is an agonist for CCR9 but is scavenged by the atypical chemokine receptor ACKR4 [25]. These examples suggest that targeting the chemokine ligand may represent an effective mechanism for remodeling immune responses.

To study the chemokine receptor:chemokine axis in more detail, a variety of chemokine modulators is needed. However, these small proteins have proven challenging to target effectively using conventional approaches. For example, an NMR-based CCL28 fragment screen gave a low hit rate (<1%) of very weak affinity binders (*K*d>1 mM) [26], thereby indicating low ligandability. We were therefore prompted to consider alternative approaches. With an avalanche of protein structural information becoming available, the use of de novo structure generation algorithms is making a revival. Recent advances in artificial intelligence-driven structure prediction have yielded highly accurate tools such as AlphaFold2 (AF2) [27]. Building on these developments, researchers have introduced de novo binder design frameworks that generate protein structures and sequences from scratch [28], including Chai-2 [29], AlphaProteo [30], and BindCraft [31]. The latter has emerged as a fully integrated, one-shot platform for binder design based only on an input target protein. BindCraft is an AF2-driven inverse-design pipeline that hallucinates binder sequences by backpropagating gradients through AlphaFold2-Multimer [32] and repredicting the binder-target complex at every iteration, enabling flexible co-optimization of sequence, structure, and interface. Designed binders are further refined using MPNNsol [33] and filtered based on AF2 confidence and Rosetta [34] physics-based metrics. The platform’s integrated workflow makes BindCraft easy to apply, lowering the barrier of entry for de novo binder design and enabling broad accessibility with a high success rate against a diverse set of targets [31,35,36].

Here, we apply BindCraft’s de novo protein design to generate modulators for CCL25. Out of the generated BindCraft designs, four miniproteins were selected, produced, isolated and characterized. All show selective binding to CCL25 and form complexes that inhibit receptor activation. Most prevent activation by impeding chemokine binding to the receptor, thus impairing all signaling and migratory responses, but one forms a G protein-biased complex with CCL25, preserving G protein recruitment while suppressing arrestin recruitment to CCR9, and selectively inhibits CCR9, but not ACKR4. These results highlight the effectiveness of targeting chemokines with de novo protein modulators as unique research tools to probe the receptor:chemokine signaling axis.

## Results

### Generation of de novo miniprotein binders targeting CCL25

In order to generate binders against CCL25, the AF2-predicted model was obtained from the AlphaFold protein database [37]. CCL25 is the largest member of the chemokine family and contains a long C-terminal extension following the classic chemokine fold. To focus the miniprotein designs on the core chemokine domain to block CRS interactions and receptor binding (**Fig. 1A, B**), we performed our design efforts using a truncated form of the chemokine. The truncated CCL25 (CCL25_ΔCT, 1-77) activated both CCR9 and ACKR4 with similar efficacy while showing slightly enhanced potency only towards CCR9 compared to the full-length ligand (**Fig. S1**).

The overall workflow is presented in **Fig. 1C**. Starting from the CCL25 AlphaFold2 model, the BindCraft algorithm generates candidate designs of 75-125 amino acids with diverse structures. No predefined binding hotspots were imposed, and thus, designs were permitted to explore the entire CCL25 surface, allowing BindCraft to identify favorable interaction sites. At most two proteins with identical backbones were used for further processing to promote diversity among the designed binders. Following the design run, after 101 candidates passed computational filters, binders were ranked by the ipTM score.

Predicted complexes demonstrated robust prediction metrics, with ipTM scores (interface predicted TM-score) ranging from 0.55 to 0.87, ipAE scores (interface predicted aligned error) from 0.16 to 0.34, and pLDDT scores (predicted local distance difference test) from 0.83 to 0.95 (**Fig. S2**). Structural inspection revealed substantial backbone diversity among designs, although all adopted predominantly α-helical secondary structures (**Fig. S3**). Pairwise TM-score analysis yielded an average of 0.41, and UMAP projection of TM-score similarities confirmed broad structural dispersion (**Fig. S3**). Despite the absence of a predefined binding hotspot, most designs target a region encompassing one face of the α-helix, the 3_10_-turn, and the 40s-loop with large structural diversity (**Fig. 1D**). As a last step, we subjected the top 20 miniprotein designs to 100 ns MD simulations to assess complex stability. Calculation of the backbone RMSD of the binders reveals good stability for all complexes, and estimation of binding free energy using the MM/GBSA method [38] furthermore predicts favorable binding complexes of all top 20 miniproteins (**Table S1**).

### De novo designed binders target CCL25 with high affinity

From the top 20 scored binders, four were selected for insertion into expression vectors for *E. coli* purification based on predicted binding energies, diversity of structural scaffolds, and binding interfaces (**Fig. 2A, Table S1**). Three of the four VU miniproteins (VUPs), VUP25101, VUP25107, and VUP25112, are predicted to interact with the C-terminal helix of the chemokine, blocking CRS1 interactions and opposite the GPCR transmembrane bundle. The fourth, VUP25111, makes unique interactions compared to the others (**Fig. S4**) and is predicted to bind along the β1 strand and 40s-loop of CCL25 without substantially impairing either CRS1 or CRS2 binding interfaces. The binding of the designed miniproteins to CCL25 was validated by fluorescence polarization spectroscopy (FP) of a fluorophore (AZ488) attached to the C-terminus of full-length CCL25. An increase in the polarization of the fluorophore emission is the result of a decrease in the tumbling rate during fluorophore excitation and is reflective of an increase in the size of the protein complex due to miniprotein binding (**Fig. 2B**). All four tested binders produced an increase in polarization, indicating that the miniproteins indeed bind the CCL25 target (**Fig. 2C**). Similar affinity values were calculated for all four VUPs (VUP25101: pK_d_ = 6.42 ± 0.74, VUP25107: pK_d_ = 6.82 ± 0.43, VUP25111: pK_d_ 6.36 ± 0.35, VUP25112: pK_d_ = 6.30 ± 0.41). Importantly, these results also confirm that the miniproteins bind the full-length CCL25 chemokine despite being designed against an artificially truncated form, CCL25_ΔCT.

**Figure 2.**
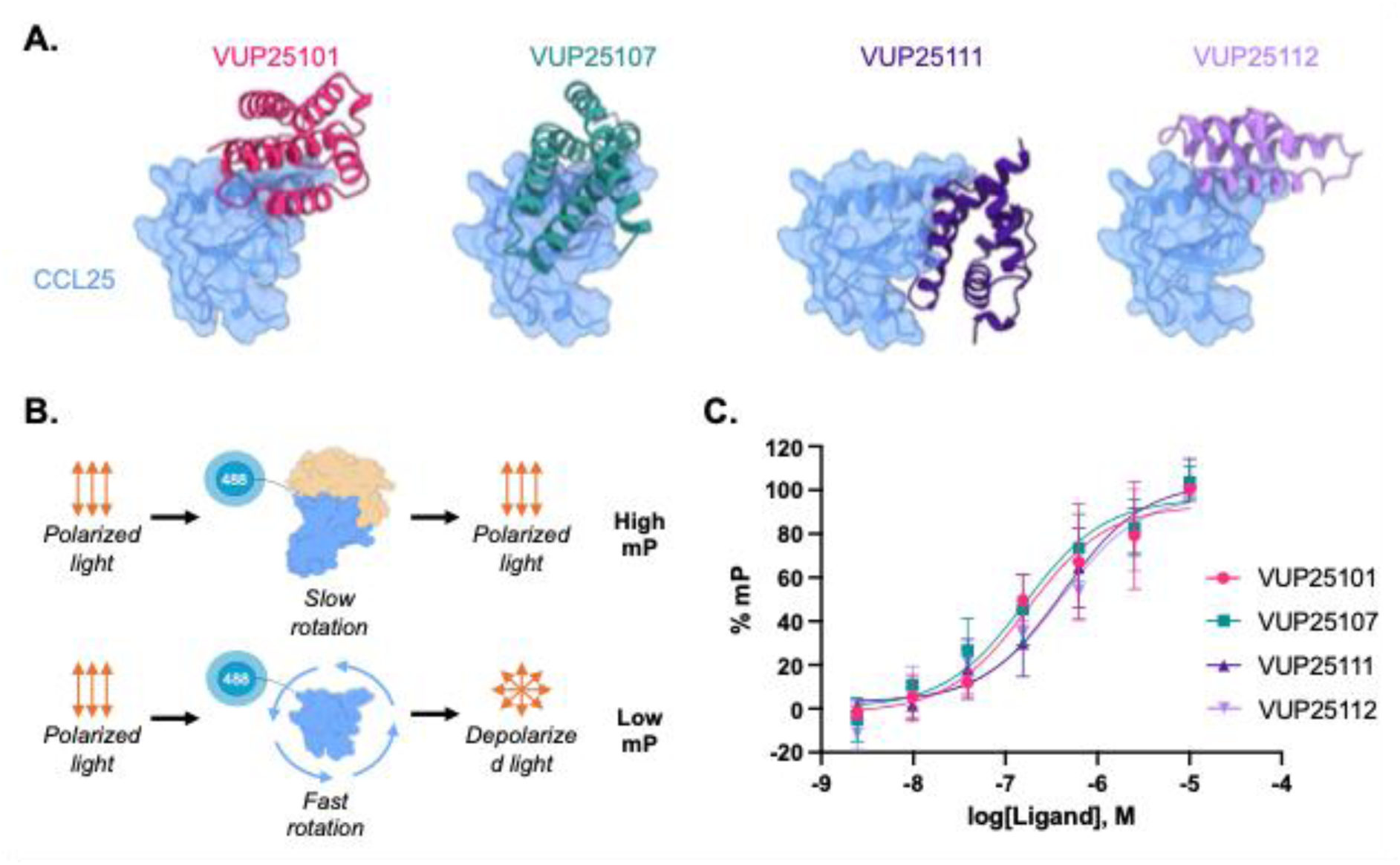
VUPs (VU miniproteins) are confirmed to directly bind CCL25. (**A**) Structural models of the generated designs for CCL25. CCL25 is shown in blue with a surface representation, and the binders are displayed as cartoon in various colors. (**B**) Cartoon schematic for the fluorescence polarization assay. When the miniprotein binds CCL25_AZ488, the size increases, as does the polarization of the emitted light. (**C**) The polarized fluorescent emission was measured for 15 nM CCL25_AZ488 in the presence of increasing concentrations of VUP binders. Binding of the miniproteins leads to an increase in polarization. Data points represent the average ± SD of three independent experiments and internally normalized to the minimum and maximum polarization of each binder.

### Miniproteins block CCL25 binding to CCR9 and ACKR4

As the miniproteins designed against CCL25 bind the desired chemokine in purified form, we next determined whether the CCL25-miniprotein complex inhibits binding to the receptor as intended. Binding of CCL25 to CCR9 or ACKR4 was determined by Time-Resolved Fluorescence Resonance Energy Transfer (TR-FRET) between a lanthanide donor attached to a SNAP-tag fused to the receptor N-terminus and a fluorophore on the C-terminus of CCL25 (AZ647) (**Fig. 3A**). CCL25_ΔCT was used in place of the full-length chemokine to shorten the distance between the donor and acceptor and improve energy transfer efficiency. Incubating CCL25 with three of the miniproteins, VUP25101, VUP25107, and VUP25112, impaired binding to CCR9 resulted in a complete loss of FRET at high miniprotein concentration (**Fig. 3B**). For ACKR4, VUP25101 and VUP25107 exhibit comparable inhibitory potencies, whereas VUP25112 is slightly reduced (**Fig. 3C**). Together, these these findings confirm that the miniproteins act as potent inhibitors by directly interfering with CCL25 binding to the receptors. VUP25111 shows a drastically lower inhibitory potency compared to the other binders and did not completely eliminate the binding signal. Since the binder readily complexes with the chemokine at these concentrations (**Fig. 2**), this suggests that the CCL25:VUP25111 complex retains some ability to interact with the receptor.

**Figure 3.**
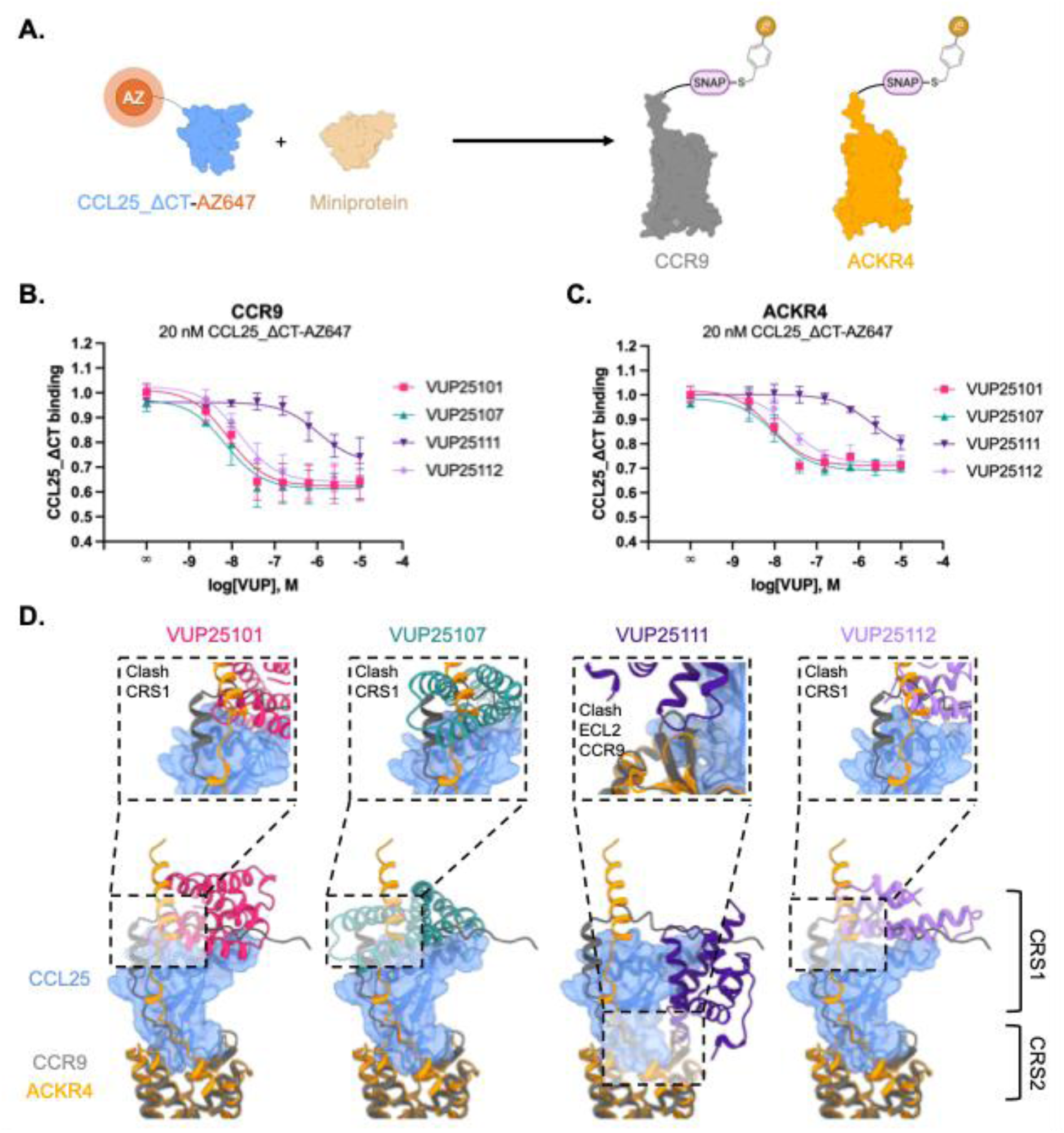
VUPs interfere with CCL25 binding to CCR9 and ACKR4 through clashes with CRS1. (**A**) Design of binding assay using TR-FRET between CCL25_ΔCT-AZ647 and terbium (Tb) labelled SNAP-CCR9 and SNAP-ACKR4. TR-FRET binding results for CCR9 (**B**) and ACKR4 (**C**) using 20 nM CCL25_ΔCT-AZ647 with increasing concentrations of the miniproteins, which shows a decreasing TR-FRET signal indicating an inability of the chemokine:VUP complex from binding the target receptor. Values represent the mean ± SD of three independent experiments performed in triplicate, presented as fold over vehicle condition. Significance was determined using Extra sum-of-squares F test with VUP25101 as a reference, *P < 0.01, **P < 0.0001. (**D**) Predicted structural models of miniproteins superimposed on the AF3-predicted CCR9:CCL25 (dark blue cartoon) and ACKR4:CCL25 (golden cartoon) complexes with zoom-in panels highlighting where the miniproteins are predicted to sterically clash with receptor:chemokine models.

To explore a possible structural explanation for how the binders prevent CCL25 from engaging its receptors, we modeled the CCR9:CCL25 and ACKR4:CCL25 complexes using AlphaFold3 [39]. Superimposing the miniprotein:CCL25 complexes onto the predicted receptor:chemokine complexes reveals steric clashes in CRS1 between VUP25101, VUP25107, and VUP25112 and the N-terminus of both CCR9 and ACKR4 (**Fig. 3D**). This suggests that disrupting solely these contacts is sufficient to block proper receptor:chemokine complex formation. In contrast, superimposing the VUP25111:CCL25 complex on the models shows only minor clashes with the extracellular loop 2 (ECL2) of CCR9 and no predicted conflicts with ACKR4. Together, these results indicate that blocking CRS1 interactions between the receptor N-terminus and chemokine body robustly prevents CCL25 binding to both its canonical and atypical receptors, while targeting the β1-strand appears structurally tolerated.

### De novo miniproteins inhibit CCL25-mediated arrestin recruitment to CCR9 and ACKR4

The effect of CCL25-targeting on receptor activation was first evaluated by assessing arrestin recruitment. Both CCR9 and ACKR4 show robust engagement with arrestins following agonist stimulation and serve as simple readouts of changes in receptor activation [40,41]. Thus, β-arrestin2 recruitment to the receptors was measured by BRET between a C-terminal luciferase fusion on the receptor (RlucII) and an N-terminal GFP10 on β-arrestin2 (**Fig. 4A**). In both cases, the addition of chemokine leads to a substantial increase in resonance energy transfer, indicating translocation of the effector in response to agonist-mediated receptor activation (**Fig. 4B, C**). Adding an increasing concentration of the miniprotein binders with a constant 20 nM CCL25 (Full-Length) progressively decreased the CCL25-promoted activation and all binders completely inhibited CCR9-arrestin recruitment at the highest concentrations (**Fig. 4B**). VUP25101, VUP25107, and VUP25112 showed similarly effective suppression of CCL25-mediated activation of CCR9 (VUP25101: pIC_50_ = 7.92 ± 0.07; VUP25107: pIC_50_ = 8.01 ± 0.24; VUP25112: pIC_50_ = 7.56 ± 0.41). The IC_50_ of VUP25111 was dramatically shifted by 2 log units (pIC_50_ = 5.77 ± 0.33). Given the different binding mode of VUP25111 to CCL25 (**Fig. 3D**), this indicates that the mechanism by which VUP25111 inhibits CCR9 activation may also differ.

**Figure 4.**
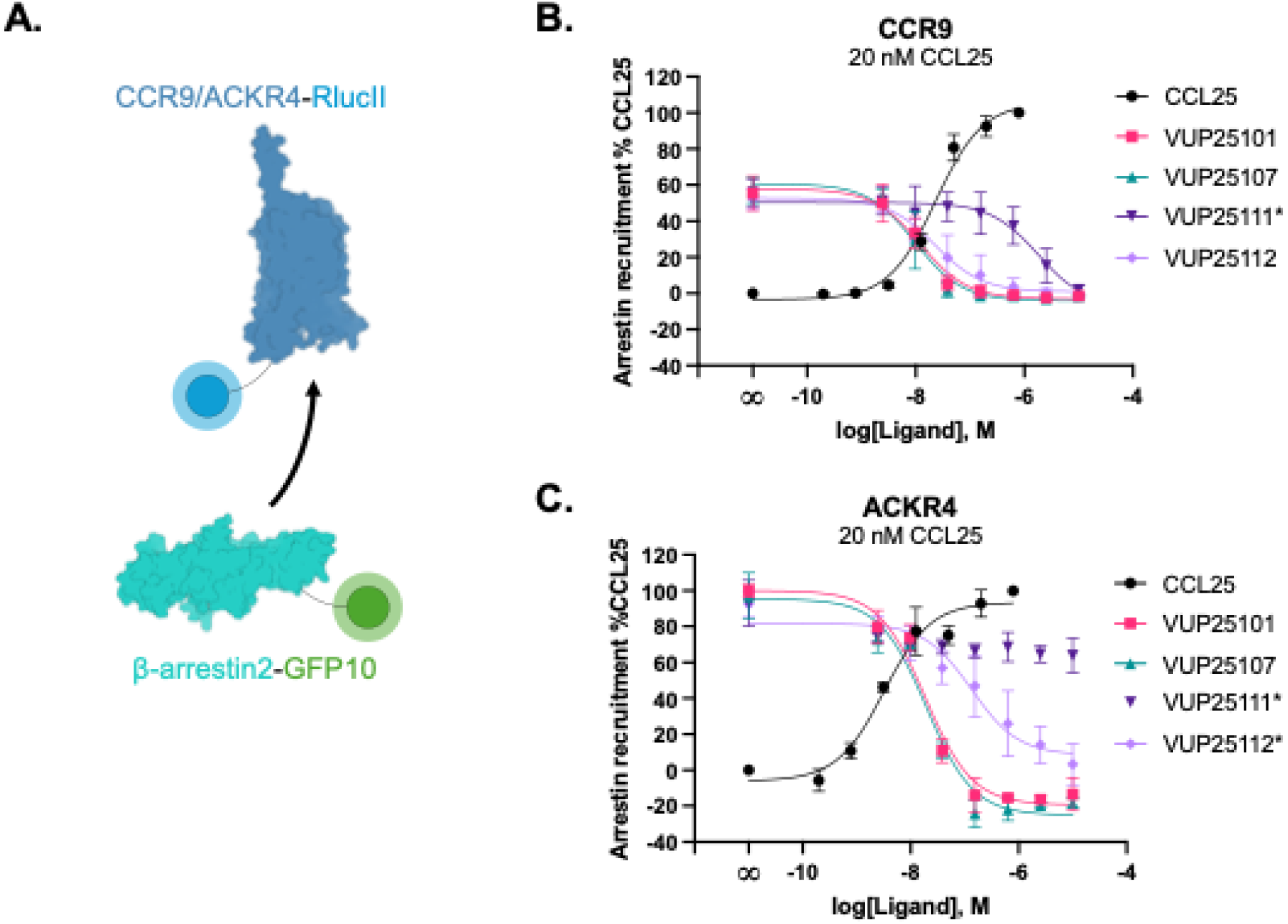
VUPs selectively inhibit CCL25-induced arrestin recruitment towards CCR9 and ACKR4. (**A**) Schematic illustration of BRET sensor pair CCR9/ACKR4-RlucII and GFP10-β-arrestin2. The structure of β-arrestin2 is generated using AF3. β-Arrestin2 recruitment towards CCR9 (**B**) or ACKR4 (**C**) following stimulation across a titration of CCL25 concentrations (black) or a fixed concentration of CCL25 (20 nM) preincubated with increasing concentration of miniprotein. Values represent the mean ± SD of three independent experiments performed in duplicate and normalized to min and max CCL25 response. Significance was determined using Extra sum-of-squares F test with VUP25101 as a reference. *P < 0.0001.

CCL25-mediated arrestin recruitment to ACKR4 was also robustly inhibited by VUP25101 (pIC_50_ = 7.73 ± 0.74) and VUP25107 (pIC_50_ = 7.71 ± 0.43) (**Fig. 4C**). VUP25112 also completely impaired CCL25-mediated activation of the atypical receptor at the highest concentrations, but with significantly less potency (pIC_50_ = 6.95 ± 0.34) compared to VUP25101 and VUP25107 (**Table S2**). The unique binder VUP25111, on the other hand, did not significantly impact the activation of ACKR4 by CCL25 at any tested miniprotein concentration. The lack of effect supports the structural interpretation that the CCL25:VUP25111 complex can bind ACKR4 as a cooperative unit and implies that this complex also does not alter the receptor’s native, chemokine-promoted activation. These data indicate that CCL25-activation of both CCR9 and ACKR4 can be efficiently inhibited by targeting the CRS1-binding interface and suggest sufficient differences in the CCL25:receptor interface that allow receptor-specific inhibition.

CCL25 specificity was experimentally evaluated by testing the miniproteins against CXCL12 by tracking arrestin recruitment to CXCR4 and ACKR3, the native agonist for this canonical and atypical receptor pair (**Fig. S5**). VUP25111 and VUP25112 showed no effect on CXCL12-mediated activation at any concentration, while VUP25101 and VUP25107 showed partial inhibition at the highest tested concentration (10 µM).

In addition, the effects of the VUPs were evaluated against CCL20, a closely related chemokine to CCL25, by assessing arrestin recruitment to CCR6 and ACKR4 (**Fig. S6**). VUP25107 and VUP25112 displayed similar inhibitory potencies of CCL20 as to CCL25 for ACKR4 (VUP25107: pIC50 = 7.72 ± 0.21; VUP25112: pIC50 = 6.93 ± 1.03). In contrast, VUP25101 showed reduced inhibition of CCL20 relative to CCL25 (VUP25101: pIC50_CCL20_ = 6.06 ± 0.13 v pIC50_CCL25_ = 7.73 ± 0.08), indicating greater selectivity for the designed target. CCL20 activation of CCR6 further supports the specificity of VUP25101, which only showed a modest effect at the highest concentration tested. VUP25111 showed no inhibitory effects on CCL20 activation of either CCR6 nor ACKR4, which may indicate a lack of binding to the chemokine, but is also indiscernible from forming a receptor competent complex similar to the proposed CCL25:VUP25111:ACKR4 interaction.

### G protein activation by CCR9 is inhibited by VUP25101, VUP25107, and VUP25112, but not VUP25111

Immune cell migration is driven in part by G protein coupling, thus, inhibition of this function is critical for crafting immune responses. The ability of the de novo binders to impact canonical G protein coupling was tested first by measuring the recruitment of ‘mini’ G proteins (mG_i_), engineered Ras domain from Gα_s_ with the interacting residues of G_i_ [42], to CCR9 by BRET between CCR9 C-terminally tagged with RlucII and mG_i_ fused to a Venus fluorophore (**Fig. 5A**). Like with arrestin recruitment, both VUP25101 and VUP25107 inhibited CCL25-induced mGi recruitment with similar potency (VUP25101: pIC_50_ = 8.17 ± 0.21; VUP25107: pIC_50_ = 8.44 ± 0.17) (**Fig. 5B**). Despite potent inhibition of arrestin engagement, VUP25112 showed a right-shifted IC_50_ value compared to the VUP25101 and VUP25107 (pIC_50_ = 7.15 ± 0.51). Interestingly, VUP25111 showed no effect on the mG_i_ recruitment even at concentrations that robustly inhibited arrestin engagement. The deviations between the arrestin results (**Fig. 4**) and G protein (**Fig. 5B**) suggest that the specific ligand interactions or receptor conformations driving arrestin engagement with CCR9 may be more strict than for the G proteins.

**Figure 5.**
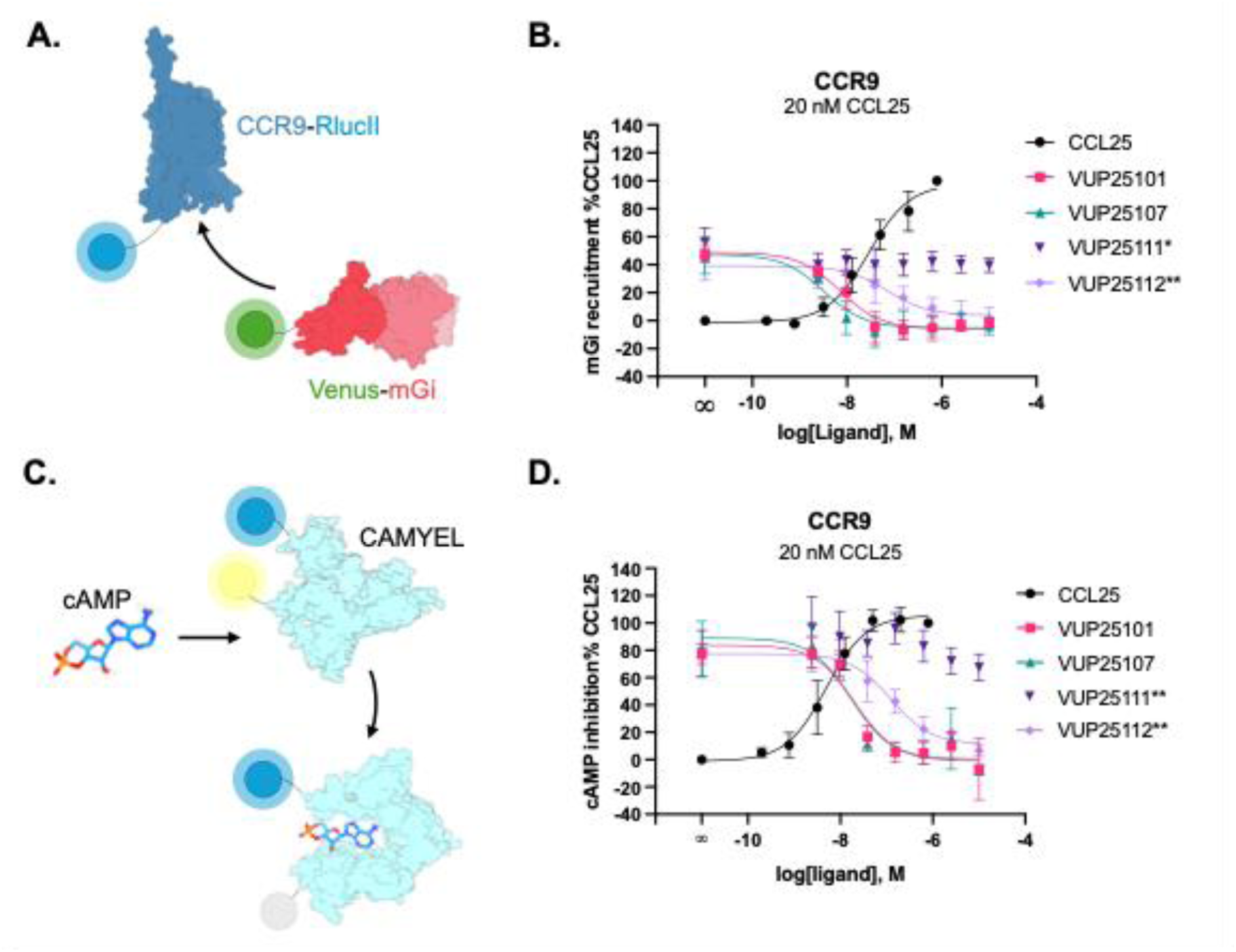
Most VUPs, but not VUP25111, effectively inhibit CCL25-induced G_i_ protein activation. (**A**) Schematic illustration of BRET pair sensor CCR9 and Venus-mG_i_. The complex representing mGi is taken from PDB 9KO4 [45]. (**B**) Concentration-dependent mG_i_ recruitment to CCR9 following stimulation with a titration of CCL25 (black) or with 20 nM CCL25 preincubated with increasing concentrations of miniproteins. (**C**) Suppression of cAMP signaling monitored by the BRET-based cAMP sensor CAMYEL. The structure of CAMYEL is generated using AF3. (**D**) CCR9 induced cAMP suppression following stimulation with a titration of CCL25 (black) or a fixed concentration of CCL25 (20 nM) preincubated with a titration of miniproteins. Values represent mean ± SD of three independent experiments performed in duplicate, normalized to min and max CCL25 response. Significance was determined using Extra sum-of-squares F test with VUP25101 as a reference, *P < 0.01 **P < 0.005.

GPCR signaling is amplified by a cascade and altered recruitment of the G protein effector may not be reflected in downstream outputs. CCR9 is G_i_-coupled and activation inhibits the production of cAMP and decreases the total concentration. The impact of the miniproteins on G protein signaling by CCR9 was monitored using a BRET-based cAMP biosensor (CAMYEL) to detect the inhibition of forskolin (FSK)-induced cAMP production (**Fig. 5C**). Downstream signaling was inhibited similar to mG_i_ recruitment, confirming the CCL25-targeting proteins are effective neutralizers of the chemokine-mediated signal (**Fig. 5D**). Despite the higher potency of CCL25 for cAMP inhibition compared to arrestin recruitment, a fixed concentration of 20 nM CCL25 was used throughout this and subsequent assays to maintain a consistent CCL25:miniprotein stoichiometry. As this concentration represents different points on the respective concentration-response curves, cross-assay comparisons of apparent miniprotein potency should be interpreted qualitatively. Like with mG_i_, VUP25111 did not impact cAMP inhibition at any concentration, further confirming that the binder is not affecting G protein activation. Taken together with the complete suppression of arrestin recruitment, it appears that the binding of VUP25111 converts the balanced natural agonist CCL25 to G protein-biased towards CCR9.

### VUP25111:CCL25 only partially impairs GPCR kinase 3 translocation to CCR9

The preserved G protein response from CCR9 (**Fig. 5**) when treated with the VUP25111:CCL25 complex suggests greater complexity to how the miniprotein suppresses arrestin interactions. Arrestins are recruited to activated CCR9, like other GPCRs, following phosphorylation of the receptor C-terminus by GPCR kinases (GRKs) [41]. Thus, translocation readouts as seen in **Fig. 4** report the combination of arrestin engagement and GRK phosphorylation and alterations to either event impact the response. To resolve if VUP25111 impairs the CCL25-mediated arrestin response through GRK phosphorylation, we measured the impact of the miniprotein binders on GRK3 recruitment to CCR9 by BRET between a nanoluciferase (Nluc) on GRK3 and a fluorophore (mV) on the receptor C-terminus (**Fig. 6A**). We have previously shown that, while all GRKs contribute of CCR9 phosphorylation, GRK3 shows the largest change in BRET upon recruitment to CCR9 [41]. Similar to G protein and arrestin, the miniproteins which interfere with CRS1 interactions, VUP25101, VUP25107, and VUP25112, also prevent chemokine-induced relocation of GRK3 to CCR9 with similar inhibitory potency as arrestin and G proteins (VUP25101: pIC_50_ = 8.40 ± 0.04; VUP25107: pIC_50_ = 8.32 ± 0.10; VUP25112: pIC_50_ = 7.81 ± 0.46) (**Fig. 6B**). Unexpectedly, VUP25111 treatment only partially inhibited GRK3 recruitment, but not the complete inhibition observed for these concentrations with arrestin recruitment. This suggests that either activation of CCR9 by the VUP25111:CCL25 complex promotes a conformation incompatible with arrestin engagement, but still accessible for GRK phosphorylation, or the reduced GRK interaction represents the formation of a phosphorylation barcode incapable of supporting arrestin recruitment. Without direct phosphorylation measurements these possibilities cannot be separated.

**Figure 6.**
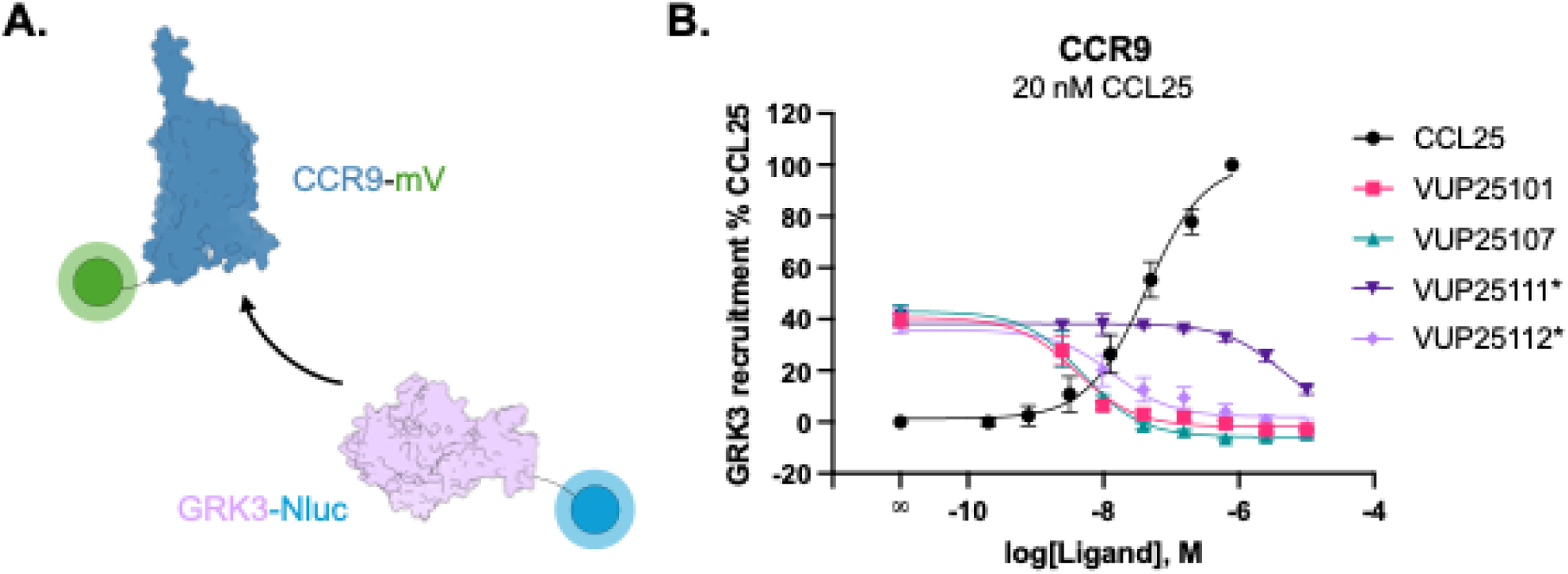
CCL25-mediated GRK3 recruitment to CCR9 is suppressed by VUP25101, VUP25107, and VUP25112, but not VUP25111. (**A**) Schematic presentation of the GRK3 recruitment BRET setup to detect GRK3-Nluc translocation to activated CCR9-mV. (**B**) Recruitment of GRK3 to CCR9 measured by BRET across a concentration range of either CCL25 (black) or VUP miniproteins with a constant 20 nM CCL25. Values represent mean ± SD of three independent experiments performed in duplicate, normalized to min and max CCL25 response. Significance was determined using Extra sum-of-squares F test with VUP25101 as a reference, *P < 0.0001.

### Targeting CCL25 with miniproteins inhibit CCR9 internalization

GRK phosphorylation largely drives CCR9 internalization through coordinating arrestins as well as a not fully-described phosphorylation dependent mechanism [41]. CCR9 internalization was tracked by BRET between CCR9-RlucII and a fluorophore (rGFP) anchored at the plasma membrane via a CAAX domain (rGFP-CAAX) (**Fig. 7A**) [43]. Receptor trafficking away from the surface is then reported as a decrease in the BRET. Binding of VUP25101, VUP25107, or VUP25112 with CCL25 potently prevented the chemokine-mediated endocytosis of CCR9 (VUP25101: pIC_50_ = 8.30 ± 0.12; VUP25107: pIC_50_ = 8.35 ± 0.16; VUP25112: pIC_50_ = 8.04 ± 0.33) (**Fig. 7B**). Internalization was also blocked fully by VUP25111, albeit at substantially lower inhibitory potency (pIC_50_ = 5.09 ± 0.27). These results suggest that the VUP25111:CCL25 complex suppresses both the arrestin dependent and independent internalization mechanisms used by CCR9, despite likely allowing for some, albeit perhaps insufficient, GRK phosphorylation.

**Figure 7.**
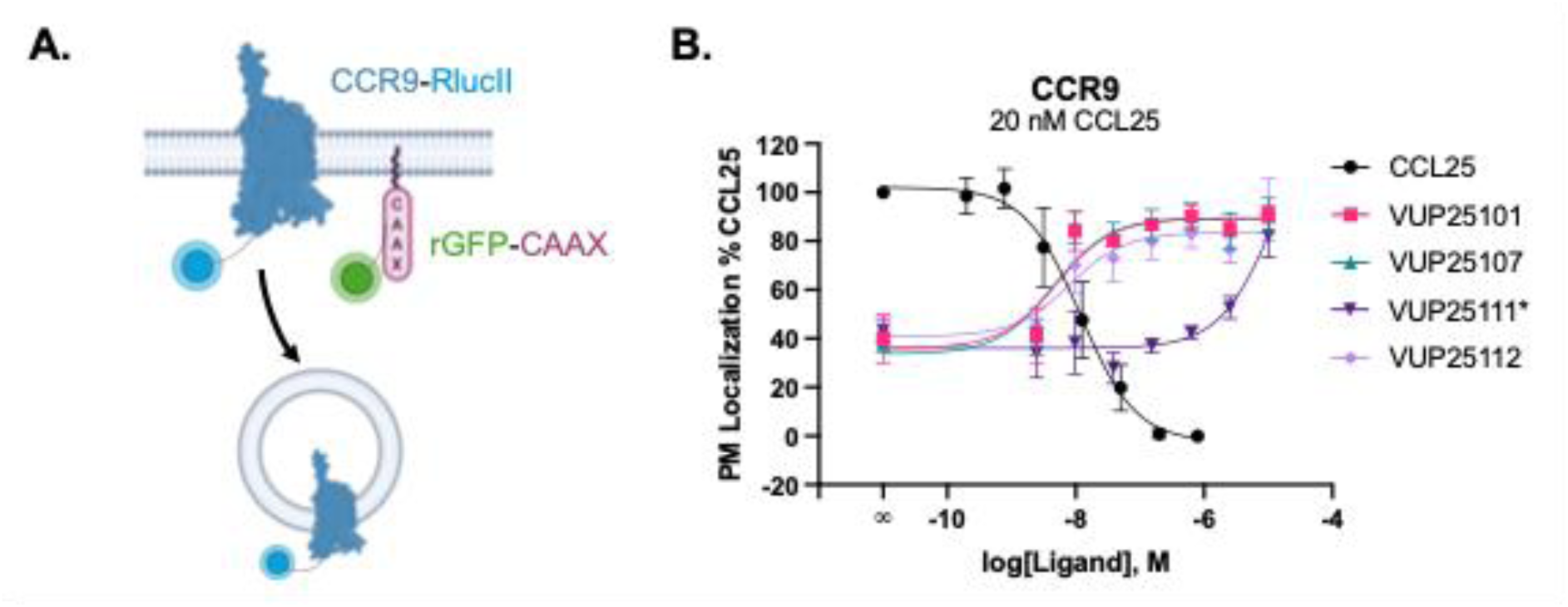
All VUPs inhibit CCL25-promoted internalization of CCR9. (**A**) Schematic representation of the BRET assay to track CCR9 internalization by monitoring the resonance energy transfer between CCR9-RlucII and rGFP-CAAX. The BRET between the probes decreases as the receptor trafficks away from the membrane through internalization. (**B**) Internalization response measured by the BRET setup presented in (**A**) across a range of concentrations of CCL25 (black) or VUP miniproteins with 20 nM CCL25. Values represent the mean ± SD of three independent experiments performed in duplicate, normalized to min and max CCL25 response. Significance was determined using Extra sum-of-squares F test with VUP25101 as a reference. *P < 0.0001.

### Immune cell migration is inhibited by targeting CCL25 at the CRS1 interface

The goal of targeting CCL25 is to shape the CCR9-driven immune response to address inflammatory diseases of the bowels and inhibit CCR9-mediated metastasis of cancer cells. Migration of MOLT-4 cells, T lymphoblasts with native CCR9 expression (lacking significant ACKR4 expression [44]), was detected in a transwell format and quantified by counting the cells in the lower chamber with flow cytometry. The CCL25 concentration corresponding to the peak migratory signal (**Fig. 8A**) was added to the lower chamber with binder and total migrated cells were counted after 2 hours. Like with the G protein assays in **Fig. 5**, VUP25101 and VUP25107 severely impaired the CCL25-mediated migration of CCR9-expressing immune cells (**Fig. 8B**). VUP25112 was less effective at inhibiting migration, as half of the migratory response towards CCL25 was still present, reflecting a similar relative level of inhibition as observed for G protein coupling. MOLT-4 cell migration was unaffected by the inclusion of VUP25111, consistent with the apparent G protein bias imparted by this binder to CCL25. These results demonstrate conclusively the effectiveness of miniproteins targeting the CRS1 interaction between CCR9 and CCL25. Such targeting impairs immune cell motility and has the potential to modify immune responses.

**Figure 8.**
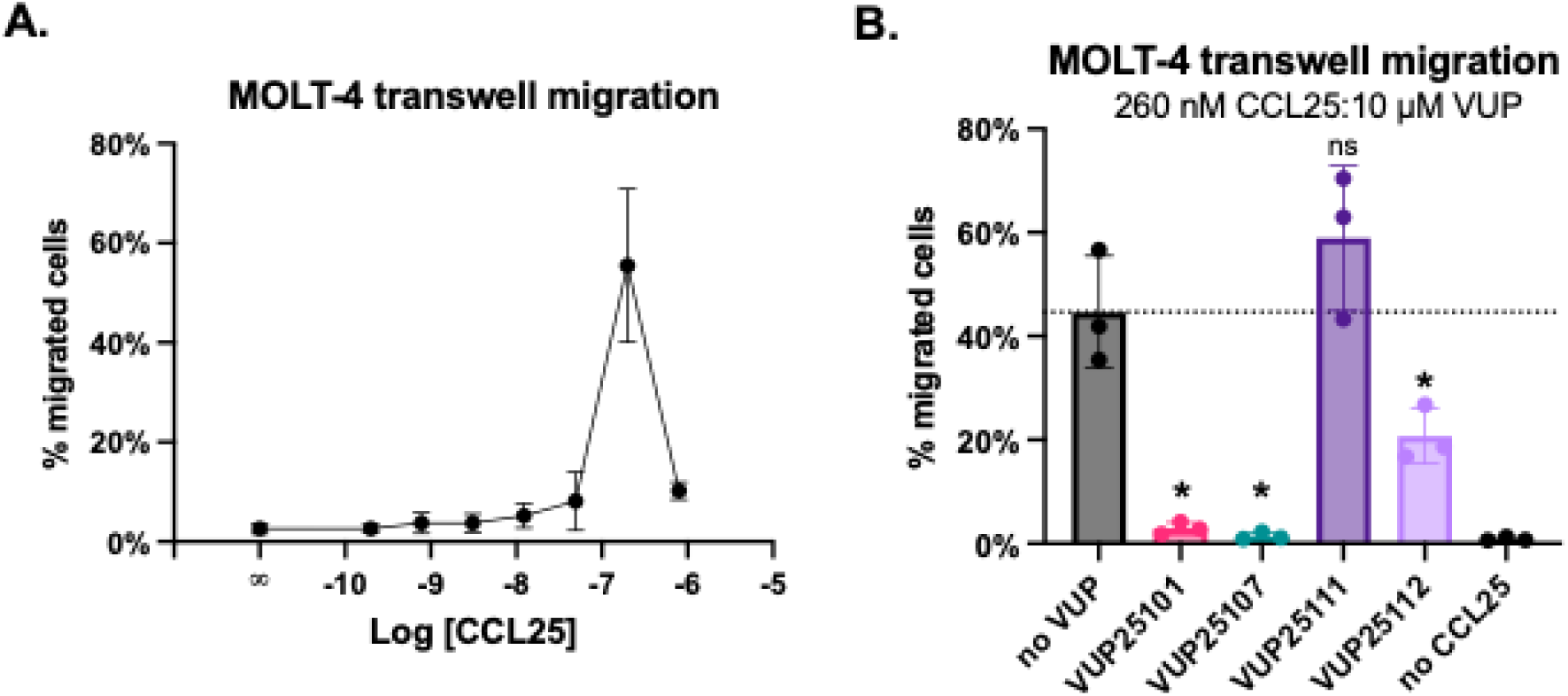
MOLT-4 chemotactic response towards CCL25 is suppressed by VUP binding. (**A**) Percentage of migrated cells towards CCL25 across a titration of chemokine concentrations. (**B**) Percentage of migrated cells towards a fixed concentration of CCL25 (260 nM, corresponding to the highest migration observed in (**A**)) preincubated with miniproteins (10 µM). Values represent the mean ± SD of three independent experiments measured in technical triplicates by flow cytometry. Significance was determined using Welch ANOVA followed by a Dunnett’s multiple comparisons test comparing to no VUP. *p<0.05.

## Discussion

Dysregulation of the CCL25:CCR9 signaling axis has been implicated in promoting inflammatory diseases of the gut [17,18] and contributing to cancer metastasis [19,20]. Although several CCR9 antagonists have been developed [46,47], none have passed clinical trials [21]. Recent advances in protein structure prediction enable accurate modeling of protein complexes and the de novo design of proteins with specific functions [48,49]. Because chemokines have proven difficult to target with small molecules, they represent an excellent application for these new protein design approaches.

Using BindCraft, miniproteins were designed to target CCL25. The miniproteins VUP25101, VUP25107, and VUP25112 were predicted to interact with the CCL25 surface involved in CRS1 interactions and robustly inhibited CCR9 and ACKR4 activation. This suggests that blocking contacts between CCL25 and the receptor N-terminus is sufficient to suppress chemokine-mediated signaling without directly targeting the receptor or its orthosteric binding pocket. Other chemokine interfaces, such as those involved in GAG binding or chemokine oligomerization, may similarly provide opportunities for modulating receptor activation. Unexpectedly, targeting CCL25 along the chemokine β1 strand, in the case of VUP25111, resulted in a miniprotein that inhibited CCR9-induced arrestin recruitment, but not G protein activation, and showed no inhibition of ACKR4. Thus, targeting CCL25 produced not only potent inhibitors, but also presented new insights into signaling along the receptor:chemokine axis (**Fig. 9**). Structural modeling suggests that when complexed with CCL25, VUP25111 would clash with the ECL2 of CCR9, whereas the three-residue-shorter loop of ACKR4 can accommodate the complex. Binding and G protein activation results suggest that VUP25111 does not prevent CCL25 from binding CCR9, but instead modifies the agonist interactions to become G protein-biased, as discussed further below. Selectively inhibiting the canonical receptor without impairing its atypical partner would suppress migratory signaling without disrupting chemokine scavenging. Blocking this regulatory mechanism may increase chemokine levels, dysregulate the signaling axis, and eventually overwhelm the inhibitors. Thus, continual removal of the agonist through native scavenging may be a necessary consideration when designing chemokine-targeting therapeutic.

**Figure 9.**
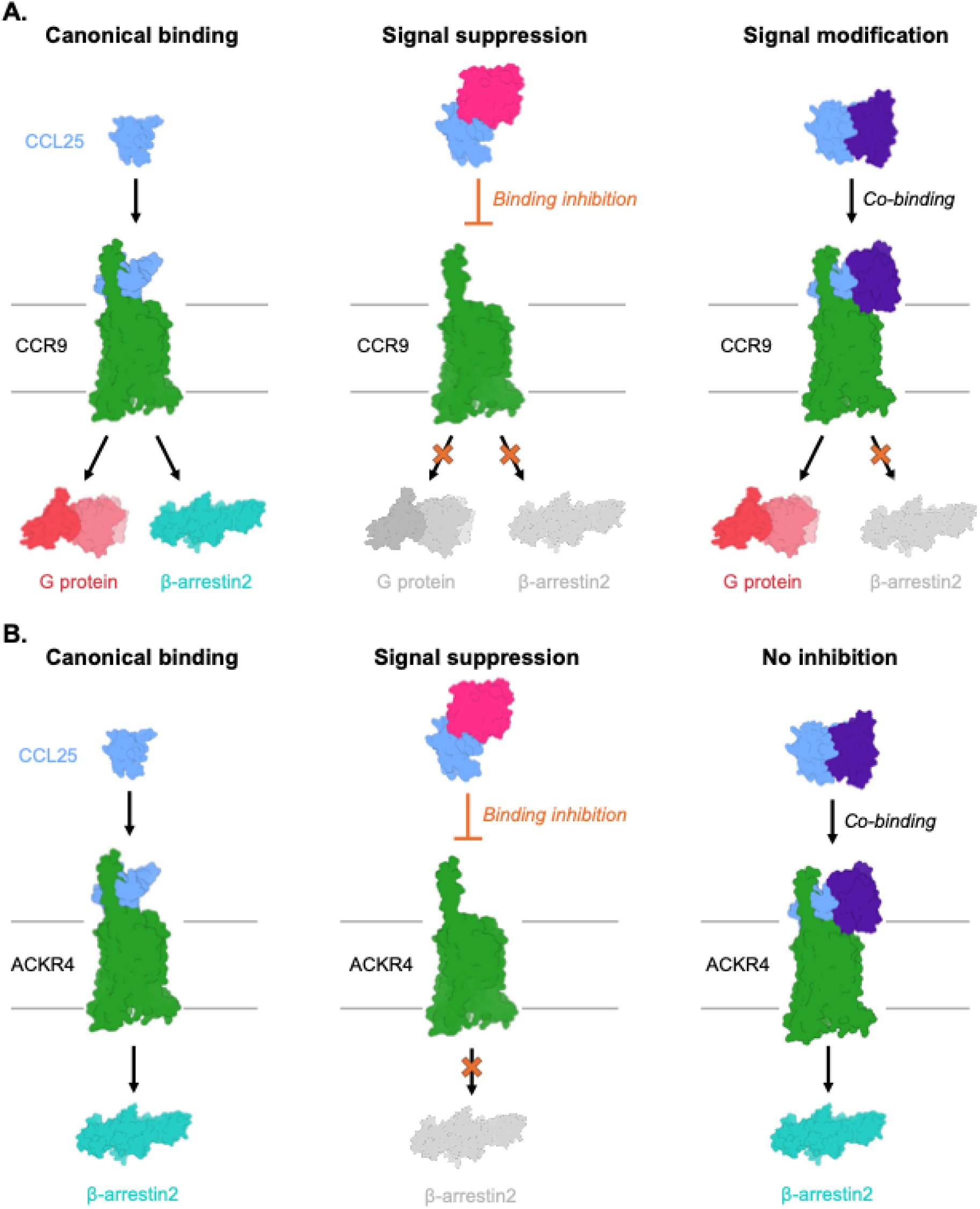
Mechanisms by which VUPs (VU miniproteins) modulate CCL25 signaling at CCR9 and ACKR4. (**A**) Canonical engagement of CCL25 with CCR9 produces a balanced signaling response, activating both G protein- and β-arrestin2-dependent pathways. Signal suppression occurs when miniproteins (VUP25101, VUP25107, and VUP25112) sterically block CCL25:CCR9 interactions, leading to reduced activation of both G protein and β-arrestin2 signaling. Signal modification can also arise from co-binding of the CCL25:miniproteins complex to the receptor (VUP25111), resulting in biased signaling characterized by selective attenuation of the β-arrestin2 pathway while preserving G protein-mediated signaling for CCR9. (**B**) Canonical engagement of CCL25 with ACKR4 results in β-arrestin2 signaling. Signal suppression occurs when miniproteins (VUP25101, VUP25107, and VUP25112) sterically block CCL25:ACKR4 interactions, leading to reduced β-arrestin2 signaling. Miniprotein VUP25111 does not inhibit activation of ACKR4 by CCL25. The complex representing G protein is taken from PDB 9KO4 [45], the structure of β-arrestin2 was generated using AF3.

Biased signaling with regard to GPCRs generally refers to preferential coupling to either G proteins or arrestins and can arise from specific ligands, receptor activation mechanisms, or cellular environment. VUP25111 prevents CCL25-induced arrestin recruitment to CCR9 but leaves G protein signaling intact. To our knowledge, this represents the first designed binder to alter the signaling bias of a natural ligand. The fact that the same ratio of binder to chemokine impairs arrestin coupling, but not G protein (**Fig S8**), indicates that the VUP25111:CCL25 complex is able to bind and activate CCR9. While GRK recruitment is partially suppressed, the lack of complete inhibition likely indicates that the kinases can phosphorylate the VUP25111:CCL25-activated receptor. The extent of phosphorylation is not clear and the inhibited GRK recruitment may reflect the formation of sub-phosphorylated states which cannot support arrestin recruitment, nor promote efficient receptor internalization. Alternatively, the miniprotein:chemokine complex may induce a conformational state incompatible with arrestin engagement. Due to the clash with ECL2, the loop would need to distort to accommodate the larger ligand complex and this displacement may translate into the G protein and GRK permissive structural state. A similar signaling profile with intact G protein but no arrestin coupling was recently observed by a CCR9 point mutation (T208^5.32^A) immediately following ECL2 [50], however, we were unable to replicate these dramatic results (**Fig. S7)**. Instead, the smaller residue showed exactly WT-like effector coupling in our experiments. The β1-binding interface of VUP25111 is similar to how CXC chemokine ligands, like CXCL12, dimerize. When the CXCL12 dimer binds its canonical receptor CXCR4, the second chemokine would also cause a large deflection of ECL2 of the receptor. Signaling of dimeric CXCL12 through CXCR4 is also biased completely towards G proteins, analogous to the effects of VUP25111 on CCL25 activation of CCR9 [51]. While G protein signaling was largely intact with dimeric CXCL12 activation of CXCR4, migration was eliminated, unlike what is shown here for CCR9. This suggests that different mechanisms of migration are likely present for the different receptors and different cell lines, and CCR9-mediated migration of MOLT-4 cells is exclusively dependent on G protein activation.

The miniproteins displayed varying degrees of chemokine selectivity among CCL25, CCL20, and CXCL12, including cross-reactivity for some designs. Given that only two additional chemokines were examined, these experiments provide an initial assessment of selectivity. As chemokine selectivity was not explicitly optimized during design, broader experimental profiling and negative-design strategies may be required to establish and improve miniprotein selectivity. Targeting chemokines is an evolutionarily efficient method to modulate immune responses. Atypical chemokine receptors scavenge chemokines to direct canonical receptor-mediated migration, whereas ticks and viruses target chemokines to suppress host immune responses. Here, this mechanism is exploited through de novo binder design, demonstrating the potential application of new AI methods to generate novel binders with new modalities. The miniproteins are valuable research tools to study chemokines and the receptor:chemokine signaling pathways and offer new options for drug discovery programs. These results provide a framework for therapeutically targeting difficult-to-modulate chemokine receptors and reveal unexpected mechanisms for signaling bias.

## Materials and Methods

Unless otherwise stated, all materials and reagents were purchased from Melford or Sigma-Aldrich. Prolume purple was purchased from Nanolight Technologies (Prolume Ltd.). All miniprotein expression plasmids were synthesized by and purchased from Genscript. Full length CCL25-AZ488, CCL25_ΔCT-AZ647 (1-77), and CCL20 were purchased from Protein Foundry.

### Generation of de novo binders

The three-dimensional structure of human CCL25 was retrieved from the AlphaFold Protein Structure Database [37]. To produce the mature form of the chemokine, the N-terminal signal sequence (M1-T23) was removed and residue numbering was reset to begin with Q1. Additionally, the C-terminus was truncated to L77 to focus the binders to the chemokine fold and yield a model of CCL25_ΔCT (1-77). Binder designs were generated with BindCraft [31] (version 1.5.0) using default_filters and the default_4stage_multimer protocol. Binders were allowed to have a length of between 75 and 125 amino acids. No hotspot residues were specified, thereby allowing the design algorithm to explore binding across the entire accessible surface of CCL25. In total, 101 designs were generated that passed computational filters and ranked using the ipTM score.

Comparison of structural similarity among binders was performed using the calculation of TM-scores using TM-align [52]. For each complex, the binder chain was extracted, and all pairwise combinations of generated binders were aligned using TM-align. To obtain a low-dimensional representation of structural relationships, TM-scores were converted to a distance matrix (1 – TM-score), and a two-dimensional embedding was generated using UMAP [53] (n_neighbors = 10, min_dist = 0.1, metric = “precomputed”).

Molecular interactions between the miniproteins designs and CCL25 were calculated with ProLIF [54] using the outputted structure from BindCraft.

### Molecular dynamics simulations

Complexes from BindCraft were prepared using the CHARMM-GUI [55] solution builder. Protonation states of residues were assigned at pH 7.4 within the CHARMM-GUI solution builder. For terminal group patching, the N-terminus was acetylated (ACE patch), and the C-terminus was methylamidated (CT3 patch). The complexes were solvated in a cubic water box with a 10 Å buffer and neutralized to 0.15 M NaCl. The CHARMM36m force-field [56] parameters were used for the proteins and ions, while the TIP3P model was used for water.

All MD simulations were performed using GROMACS 2025.1 [57]. Systems were first minimized using a 5000-step steepest-descent minimization (≤5000 steps; convergence threshold 1000 kJ/mol/nm). Subsequently, the system was equilibrated according to the protocol generated by CHARMM-GUI. Systems were subjected to a 125-ps NVT run with a 1 fs time step and positional restraints on all protein heavy atoms (force constants: 400 kJ/mol/nm² for backbone atoms and 40 kJ/mol/nm² for side chains). Initial velocities were generated from a Maxwell-Boltzmann distribution at 310.15 K. Temperature was maintained at 310.15 K using the v-rescale thermostat [58]. Production simulations were run for 100 ns in the NPT ensemble. The v-rescale thermostat was used to maintain a temperature of 310.15 K, and the pressure was kept at 1 bar via semi-isotropic coupling to the c-rescale barostat [59] (τp = 5.0 ps, compressibility = 4.5*10-5 bar). Bond lengths of hydrogen covalent bonds were constrained using the LINCS algorithm. Long-range electrostatic interactions were calculated with the particle mesh Ewald method and the Verlet cutoff scheme with a 1.2 nm cutoff for both PME electrostatics and Lennard-Jones interactions, the latter using a force-switch modifier between 1.0 and 1.2 nm. Atomic positions were saved every 50 ps.

Trajectory analysis was performed using GROMACS and MDAnalysis [60]. For each system, backbone RMSD, binding-site RMSD, and all-atom RMSD were calculated. These were done with proper masking in MDAnalysis. Binding free energies were estimated using gmx_MMPBSA [61] with the MM-GBSA method (igb = 5, 0.15 M salt). For each trajectory, 1001 frames (sampled every 100 ps over a 100 ns simulation) were included.

### AlphaFold3 structure prediction

The AlphaFold3 [62] server was used to predict the structures of CCL25 (CCL25-CT, 1-77) bound to CCR9 and ACKR4. Sequences of CCL25 (UniProt: O15444), CCR9 (UniProt: P51686), and ACKR4 (UniProt: Q9NPB9) were retrieved from UniProt [63]. For each complex, five models were generated using default settings, and the top-ranked model was selected for visualization.

### DNA Constructs and site-directed mutagenesis

Human CCR9 (1-369) and ACKR4 (1-350) were cloned into a pcDNA3.1 expression vector either alone or followed by a C-terminal RlucII tag as described previously [41]. For receptor binding assays, hCCR9 and hACKR4 were cloned into a pcDNA5 expression vector, wherein the mammalian promotor was replaced with EF1a promotor, preceded by a N-terminal SNAP-tag. GFP10-β-arrestin2 (gifted from N. Heveker, Université de Montréal) [64], NES-Venus-mGs/i143 (mG_i_) [42], and mCitrine-EPAC-Rluc (CAMYEL) [65,66] were described previously. Site-directed mutagenesis was performed using the Q5 mutation kit (NEB) and confirmed by sanger sequencing.

### Chemokine purification from E. Coli

The chemokines CCL25 and CXCL12 were expressed and purified in *E.coli* following previously established protocols [41,67]. Briefly, codon optimized chemokine sequences containing the 8x histidine-tag and enterokinase cleavage site were cloned into a pET21 vector. The synthesized plasmid was transformed into *E.coli* BL21(DE3)pLysS cells and protein expression was triggered by Isopropyl-β-D-thiogalactopyranoside (IPTG) induction. Inclusion bodies containing the chemokines were harvested by sonication and solubilized in a buffer composed of 50 mM Tris, 6 M guanidine-HCl, and 300 mM NaCl at pH 8.0. Next, the solubilized protein was purified using a nickel-nitrotriacetic acid (Ni-NTA) affinity column, subsequently washed with 50 mM MES, 6 M guanidine-HCl, 300 mM NaCl pH 6.0 and eluted with 50 mM acetate, 6 M guanidine-HCl, 300 mM NaCl pH 4.0. The elution fraction was refolded in 50 mM Tris, 700 mM arginine-HCl, 1 mM EDTA, 1 mM GSSG, 200 mM glutamine, 0.1% Triton-X (v/v) pH 7.5 by dropwise addition. After refolding, the protein was dialyzed in 20 mM Tris with 150 mM NaCl pH 8.0. The 8xHis-tag was removed using Enterokinase (New England Biolabs), and cleavage was confirmed by SDS-PAGE and LC-MS. The tag-free chemokine was further purified on a Ni-NTA column, washing with 50 mM Tris pH 8.0, and elution in 6 M guanidine-HCl using either 50 mM MES (pH 6.0) or 50 mM acetate (pH 4.0) in two separate fractions. Next, these fractions were purified by reverse-phage HPLC on a Gemini C18 110A column (Phenomenex) with a linear 5-95% acetonitrile gradient in 0.1% TFA (v/v). The final purified chemokines were validated by LC-MS, lyophilized, and stored at -80 °C.

### VUP purification from E. Coli

Plasmids encoding VUP constructs, N-terminally preceded by a 6x histidine purification tag, were transformed into *E. coli* BL21(DE3)pLysS by standard heat-shock transformation and plated on LB agar containing kanamycin (100 ug/mL). A single colony of each construct was inoculated into 5-10 mL LB supplemented with kanamycin (100 ug/mL) and grown overnight at 37 °C while shaking (300 rpm) to generate starter culture. The following day, 1:25 dilutions of overnight culture were made into fresh LB containing kanamycin (100 ug/mL) and grown at 37 °C while shaking (300 rpm) until mid-log phase (OD600 ∼ 0.6-0.8). Protein expression was induced by addition of IPTG to a final concentration of 1 mM, followed by incubation of cultures for 5-6 hour at 37 °C while shaking. Cell pellets were resuspended in TBS (50 mM Tris pH 7.4, 150 mM NaCl) supplemented with DNAseI (Thermo Fisher) and protease inhibitors (cOmplete, EDTA-free) and subsequently lysed by sonication. Centrifugation was repeated under the same conditions as described earlier. Imidazole was added to the supernatant for final concentration of 20 mM. Prior to applying the supernatant, Ni-NTA columns were equilibrated in TBS with 20 mM imidazole. The columns were washed with 5 column volumes of TBS containing 20 mM imidazole to remove non-specifically bound proteins. Lastly, VUPs were eluted with TBS containing 300 mM imidazole. All fractions were analyzed by SDS-PAGE to assess the purity of the protein. Eluted VUP fractions were concentrated using a 10 kDa spin filter and buffer-exchanged with 4 column volumes of TBS and stored at -80 °C. The isoelectric point of VUP25111 is near physiological pH, thus the protein was buffer exchanged to 50 mM MES pH 6.0 150 mM NaCl to improve solubility and stability for long term storage.

### Chemokine-miniprotein complex formation by fluorescence polarization

Fluorescently tagged CCL25 (CCL25-AZ488; final concentration 15 nM) was mixed 1:1 with VUP binders prepared in TRIS buffer (50 mM, pH 7.7) supplemented with 0.05% Pluronic Acid (w/v) and dispensed into a black 384-well microplate (Greiner, F-bottom). VUP binder solutions began at 10 μM and were tested in a fourfold serial dilution series. Fluorescence polarization measurements were performed using a PHERAstar plate reader set at 25 °C. The fluorescently labeled chemokine CCL25-AZ488 was excited at 485 nm, and emission was detected at 520 nm. The target milli polarization (mP) value was set to 20 and calibrated using the positive control (CCL25-AZ488 (15 nM) in buffer alone). Polarization data was calculated according to the standard equation: 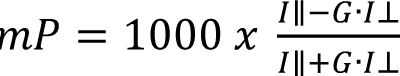

Where *I*_∥_ and *I*_⊥_ represent the fluorescence intensities measured parallel and perpendicular to the plane of polarized excitation light, respectively. The results of three independent assays were normalized to vehicle and highest response, as displayed.

### Transfection of HEK293 Cells

HEK293T cells (ATCC) were cultured in Dulbecco’s Modified Eagle’s Medium (DMEM, Thermo Fisher Scientific) containing 1% penicillin (v/v) + streptomycin (Gibco), and 10% Fetal Bovine Serum (v/v) (FBS, Bodinco) kept at 37 °C with 5% CO₂ in a humidified environment. Cells were transfected with 2 µg DNA per 1 million cells using 6 ug polyethyleneimine (PEI, Polysciences Inc.) in 150 mM NaCl. When less than 2 µg of plasmid DNA was used, the total DNA amount was adjusted to 2 µg with empty pcDNA3.1 vector. The DNA-PEI mixtures were incubated for 15 minutes at room temperature (RT). Cells were detached using trypsin/EDTA solution (Thermo Fischer Scientific) and resuspended in DMEM. Cells were counted and then added to the DNA-PEI mixture. Cells were seeded at 30k/well on a white 96-well plate (Greiner) and incubated for 48 hour before readout.

### Inhibition of CCL25 binding to receptors by TR-FRET

To make SNAP-CCR9 or SNAP-ACKR4 membranes, HEK293T cells were transfected with 2 µg of SNAP-GPCR DNA per 2 million cells using OptiMEM (Gibco) and PEI as described above. After 48 hour, cells were harvested in PBS at 4 °C and subsequently centrifuged for 10 minutes at 1500 x g. Pelleted membranes were resuspended in 80 mM NaPO_4_ + 0.05% Pluronic acid (membrane buffer) and homogenized by sonication. Next, cells were centrifuged at 22000 x g and resuspended in membrane buffer and labelled with 100 nM Tag-lite SNAP-Lumi4-Tb labeling reagent (Revvity) for 1 hour at 37 °C. To wash of free label, membranes were washed at least five times with membrane buffer. Membranes were homogenized again by sonication, snap frozen in liquid nitrogen, and stored at -80 °C. CCL25_ΔCT-AZ647 (20 nM) was incubated for 30 min at RT with increasing concentrations of the miniproteins. Next, the receptor suspension was homogenized by sonication before adding it to the readout plate (96 well half-area microplate, Greiner). The CCL25_ΔCT-AZ647+VUP mixture was added to the membranes and centrifuged at 150 x g for 3 min before incubation for 30 min at RT. TR-FRET ratio was quantified using a PHERAstar plate reader equipped with an optic module that excites at 337 nm and measures at 620 nm (Tb) and 665 nm (FRET signal) using 400 flashes per well. TR-FRET ratio was determined by dividing the FRET intensity by the Tb intensity using a ratio factor of 10000.

### β-Arrestin2 recruitment by BRET

HEK293T cells were transfected with 50 ng receptor-RlucII and 1 µg GFP10-β-arrestin2 as described previously [41]. After 48 hour incubation, cells were rinsed once with PBS and kept in Hank’s balanced salt solution (HBSS) supplemented with 0.1% bovine serum albumin (BSA; Fraction V; PanReac Applichem, w/v) (assay buffer). Next, 5 µM of Prolume Purple (Prolume Ltd.) was added and incubated for 5 minutes at 37 °C. Cells were then stimulated with increasing concentrations of either CCL25 or VUP binders that have been preincubated with 20 nM CCL25 for 30 min. A PHERAstar plate reader (BMG) was used to measure bioluminescence at 410-80 nm and 515-30 nm at 37 °C. Prior to generating dose-response curves, a kinetic response was measured to determine the time point corresponding to the maximal CCL25-induced response. Based on these measurements, a 5 min stimulation time was selected for CCR9 and a 40 min stimulation time for ACKR4. BRET values were determined as the ratio of red and blue luminescence (515-30/410-80). The results of three independent assays were normalized to the mock (no chemokine) and highest CCL25 concentration.

### mG_i_ recruitment by BRET

HEK293T cells were transfected with 50 ng CCR9-RlucII and 250 ng Venus-mG_i_ as described previously [41]. After 48 hour incubation, cells were rinsed once with PBS and kept in assay buffer. Next, 5 µM of Coelenterazine-h (CTZ-h, Promega) was added to the cells and incubated for 5 minutes at 37 °C. Cells were then stimulated with increasing concentrations of CCL25 or VUP binders that have been preincubated with 20 nM CCL25 for 30 min. Luminescence was measured using a PHERAstar plate reader using the 475-30 nm and 535-30 nm filters. BRET values were determined as the ratio of red and blue luminescence (535-30/475-80) 20 min after CCL25 addition. The results of three independent assays were normalized to the mock (no chemokine) and highest CCL25 concentration.

### cAMP measurement by BRET

HEK293T cells were transfected with 400 ng untagged CCR9 and 800 ng CAMYEL (cAMP sensor using YFP-Epac-Rluc) as described previously [41]. After 48 hour incubation, cells were rinsed once with PBS and kept in assay buffer. Next, 5 µM of Coelenterazine-h (CTZ-h, Prolume Ltd) was added to the cells and incubated for 5 minutes at 37 °C as described before. After, cAMP levels were amplified with 10 µM forskolin (FSK, Sigma-Aldrich) for 10 min. Cells were then stimulated with increasing concentrations of CCL25 or VUP binders that have been preincubated with 20 nM CCL25 for 30 min. Suppression of cAMP production was quantified by the ratio of red and blue luminescence (535-30/475-80) 5 min after CCL25 addition. The results of three independent assays were normalized to the mock (no chemokine) and highest CCL25 concentration.

### GRK3 recruitment by BRET

HEK293T cells were transfected with 250 ng CCR9-mV and 50 ng GRK3-nLuc as described previously [41]. After 48 hour incubation, cells were rinsed once with PBS and kept in assay buffer. Next, 20 µM of CTZ-400A (Bld. Pharma.) was added and incubated for 5 minutes at 37 °C. Cells were then stimulated with increasing concentrations of either CCL25 or VUP binders that have been preincubated with 20 nM CCL25 for 30 min. A PHERAstar plate reader (BMG) was used to measure bioluminescence at 475-30 nm and 535-30 nm. BRET values were determined as the ratio of red and blue luminescence (535-30/475-80) 20 min after CCL25 addition. The results of three independent assays were normalized to the mock (no chemokine) and highest CCL25 concentration.

### CCL25-mediated internalization by BRET

HEK293T cells were transfected with 50 ng CCR9-RlucII and 200 ng rGFP-CAAX as described previously [41]. After 48 hour incubation, cells were rinsed once with PBS and kept in assay buffer. Next, 5 µM of Prolume Purple (Prolume Ltd.) was added and incubated for 5 minutes at 37 °C. Cells were then stimulated with increasing concentrations of either CCL25 or VUP binders that have been preincubated with 20 nM CCL25 for 30 min. A PHERAstar plate reader (BMG) was used to measure bioluminescence at 410-80 nm and 515-30 nm. BRET values were determined as the ratio of red and blue luminescence (515-30/410-80) 20 min after CCL25 addition. The results of three independent assays were normalized to the mock (no chemokine) and highest CCL25 concentration.

### Molt-4 cell migration quantified by flow cytometry

Transwell chemotaxis assays were performed using TC inserts for 24-well plates, 8 µm pore size (Sarstedt). MOLT-4 cells (ATCC) were collected by centrifugation at 125 x g for 5 min and subsequently resuspended in RPMI 1640 + 10% FBS (Gibco, v/v) at a density of 400k/ml. Next, 80k cells were added to the inserts and placed in the 24-well plate with either an increasing concentration of CCL25 or a fixed concentration of CCL25 (260 nM) together with 10 µM of miniprotein in the bottom chamber (pre-incubated for 30 min). The plate was then incubated for 2 hour at 37°C in a humidified atmosphere with 5% CO_2_. After incubation, cells in bottom chamber were counted using a Guava easyCyte flow cytometer (Cytek) as three technical replicates. Data is presented as percentage of migrated cells relative to the initial number of cells in the insert.

### Statisical analysis

All statistical analyses were performed using GraphPad Prism 10. Information regarding graphic representation, including the use of symbols, bar plots, and error bars, is provided in the corresponding figure legends. Unless otherwise indicated, scatter plots depict the mean values obtained from three independent experiments, each performed in either duplicate or triplicate. The statistical analysis of scatterplots was done using the extra sum-of-squares F test in Prism in reference to VUP25101. In bar plots, columns represent the overall mean across independent experiments, whereas the individual data points correspond to the mean value of each separate experiment. Error bars indicate the standard deviation (SD). The statistical significance was assessed using Welch ANOVA followed by a Dunnett’s T3 multiple comparisons test comparing to no VUP.

## Funding and additional information

This publication is part of the project TRANSLATION with file number OCENW.M.24.006 which is partly financed by the Dutch Research Council (NWO) under the grant [grant ID: https://doi.org/10.61686/YKUZU08217].

