## Supplemental Information for "Selective suppression and biasing of chemokine receptors CCR9 and ACKR4 through targeting CCL25 with de novo miniproteins"

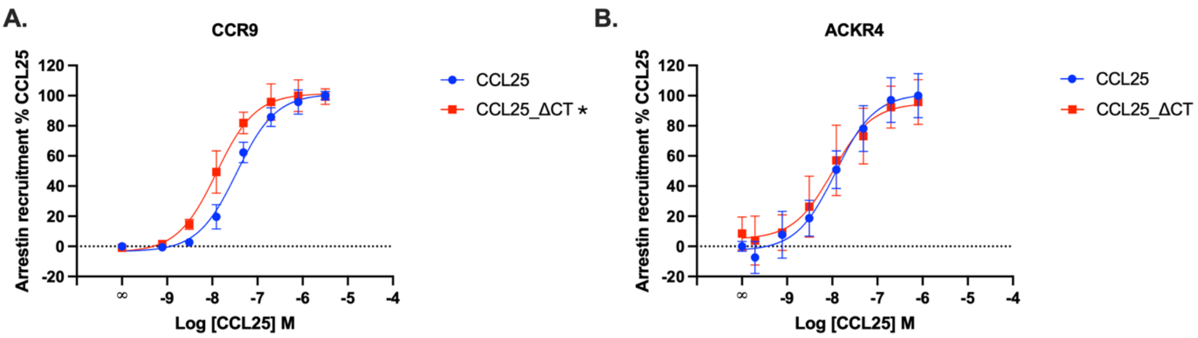


**Fig. S1. CCL25_ΔCT shows comparable efficacy, but slightly enhanced potency for CCR9.** β-Arrestin2 recruitment towards CCR9 (**A**) or ACKR4 (**B**) following stimulation across a titration of either CCL25 (blue) or CCL25_ΔCT (red) concentrations. Values represent mean ± SD of three independent experiments performed in duplicate, normalized to min and max CCL25 response. Significance was determined using Extra sum-of-squares F, *$P < 0.0001$.

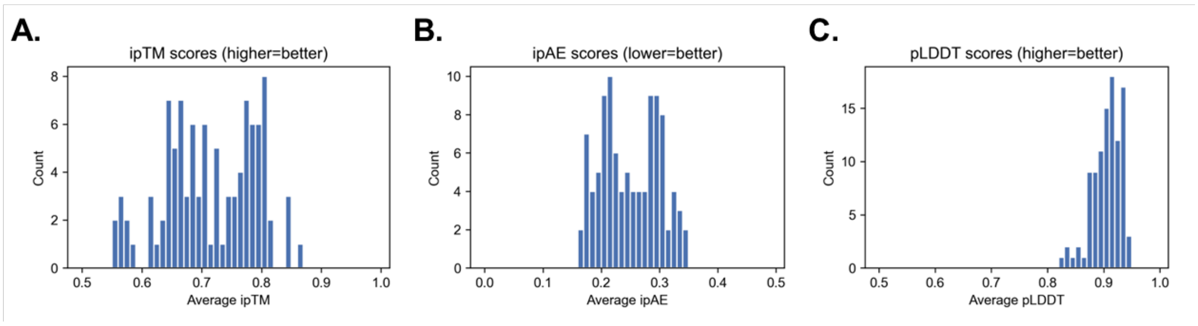


**Fig. S2. Structural confidence metrics of the 101 CCL25-targeting miniproteins generated with BindCraft.** (**A**) Distribution of interface predicted TM-score (ipTM). (**B)** Distribution of interface predicted alignment error (ipAE). (**C**) Distribution of predicted local distance difference test (pLDDT) scores.

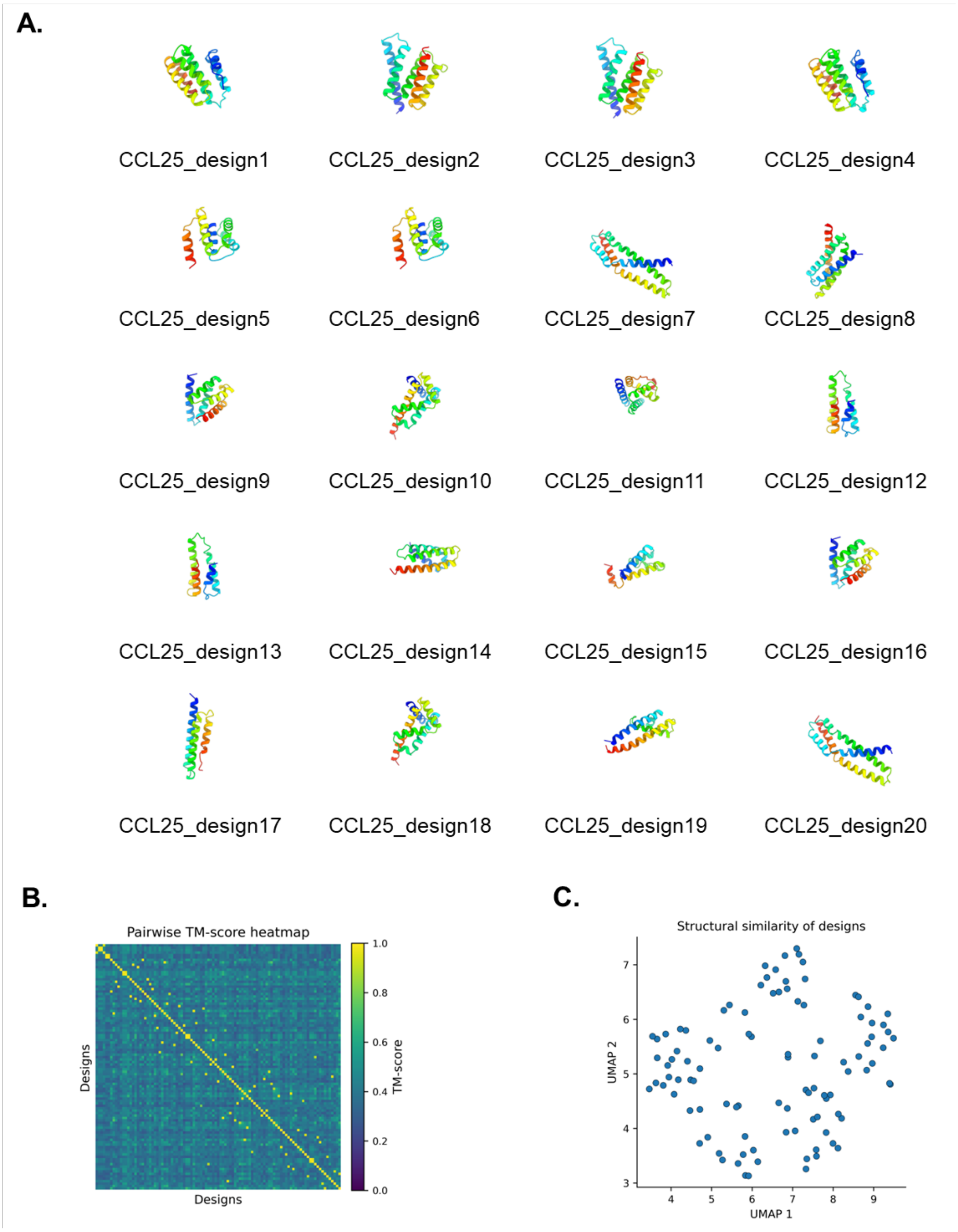


**Fig. S3. Backbone diversity of generated miniproteins.** (**A**) 3D models of the top 20 generated miniproteins against CCL25. (**B**) Matrix of pairwise TM-scores of all 101 generated miniproteins against CCL25. (**C**) UMAP projection of pairwise TM-scores.

**Table S1. Root-mean-square deviations (RMSDs) of the miniproteins and CCL25 over a 100 ns molecular dynamics simulation.** Molecular Mechanics/Generalized Born Surface Area (MM/GBSA) binding free energies, calculated from the same 100 ns MD trajectories of the CCL25:miniproteins complexes.

| **Design** | **Backbone $RMSD_{Miniprotein}$ (Å)** | **Backbone $RMSD_{CCL25}$ (Å)** | **MM/GBSA Binding free energy (kcal/mol)** |
|---|---|---|---|
| CCL25_design1 (VUP25101) | 1.46 | 2.43 | -154.21 |
| CCL25_design2 | 1.28 | 1.86 | -121.66 |
| CCL25_design3 | 1.22 | 2.61 | -110.62 |
| CCL25_design4 | 1.55 | 2.07 | -126.20 |
| CCL25_design5 | 1.73 | 2.99 | -118.33 |
| CCL25_design6 | 1.54 | 2.34 | -135.52 |
| CCL25_design7 (VUP25107) | 2.17 | 3.11 | -154.84 |
| CCL25_design8 | 2.43 | 2.87 | -107.71 |
| CCL25_design9 | 1.17 | 2.23 | -99.17 |
| CCL25_design10 | 1.20 | 2.71 | -110.85 |
| CCL25_design11 (VUP25111) | 1.97 | 1.73 | -122.46 |
| CCL25_design12 (VUP25112) | 1.93 | 2.86 | -132.45 |
| CCL25_design13 | 1.86 | 2.50 | -119.73 |
| CCL25_design14 | 1.36 | 1.73 | -105.71 |
| CCL25_design15 | 2.46 | 1.83 | -109.89 |
| CCL25_design16 | 1.33 | 2.32 | -101.75 |
| CCL25_design17 | 1.24 | 2.46 | -93.56 |
| CCL25_design18 | 1.28 | 2.22 | -113.04 |
| CCL25_design19 | 2.14 | 3.78 | -113.48 |
| CCL25_design20 | 1.70 | 2.82 | -123.46 |

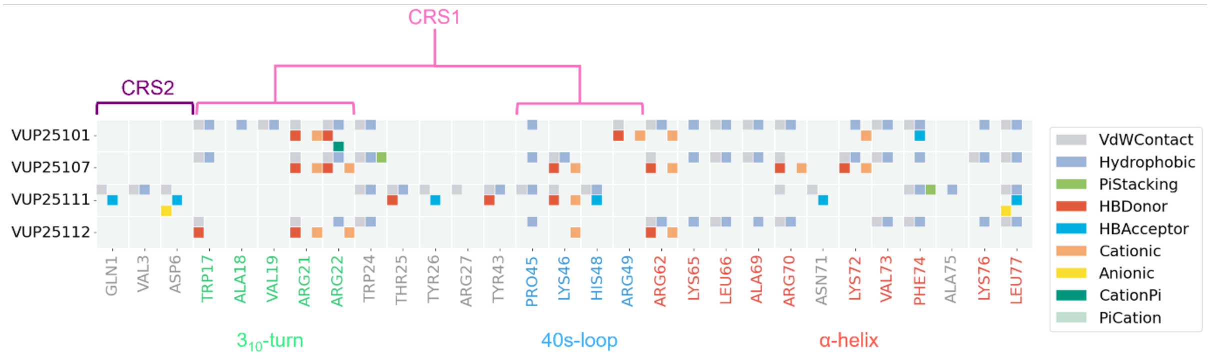


**Fig. S4. Interaction fingerprints of selected binders.** Molecular interactions of experimentally tested miniproteins based on the predicted miniproteins:CCL25 complexes as outputted by BindCraft. Residues part of CRS1 and CRS2 are highlighted in pink and purple, respectively. CCL25 residues are coloured to highlight the most frequently targeted structural elements as shown in Fig 1D: the α-helix, the 3$_{10}$-turn, and the 40s-loop, which are highlighted in red, green, and blue, respectively.

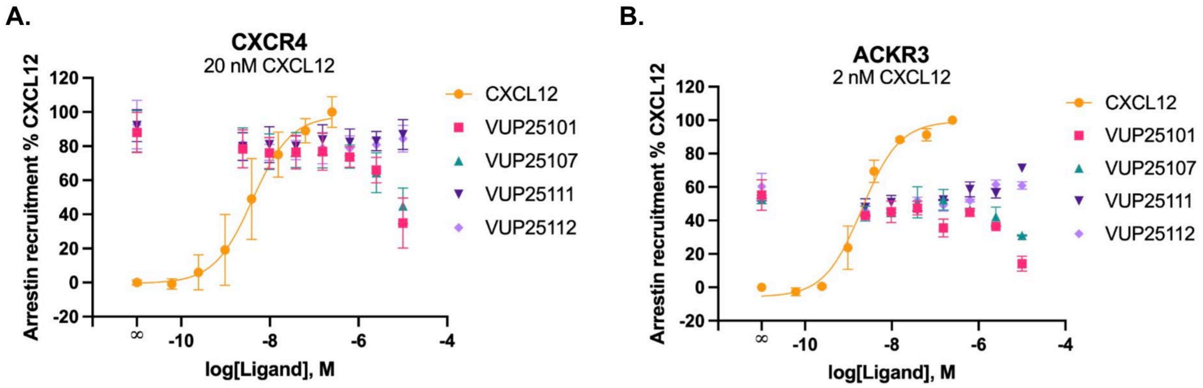


**Fig. S5. Some CCL25-targeting miniproteins impact the activation of CXCR4 and ACKR4 by CXCL12, but only at the highest concentrations.** β-Arrestin2 recruitment towards CXCR4 (A) or ACKR3 (B) following stimulation across a titration of either CXCL12 (yellow) concentrations or a fixed concentration of CXCL12 preincubated with increasing concentration of miniprotein. Values represent mean ± SD of three independent experiments performed in duplicate, normalized to min and max CCL25 response.

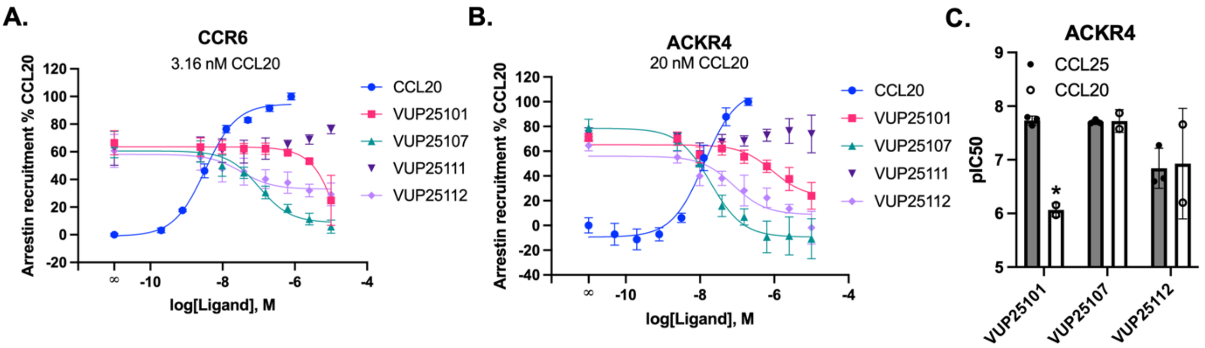


**Fig. S6. VUP25101 is selective for CCL25 over CCL20, while other miniproteins show similar effects for both chemokines.** β-Arrestin2 recruitment towards CCR6 (A) or ACKR4 (B) following stimulation across a titration of either CCL20 (blue) concentrations or a fixed concentration of CCL20 preincubated with increasing concentrations of miniprotein. (C) Comparison of the calculated IC50 values for the miniproteins for suppressing the activation of ACKR4 by CCL25 (from Fig. 4) or CCL20. Values represent mean ± SD of at least two independent experiments performed in duplicate, normalized to min and max CCL20 response. Statistical significance was determined by an unpaired t test. *P < 0.001.

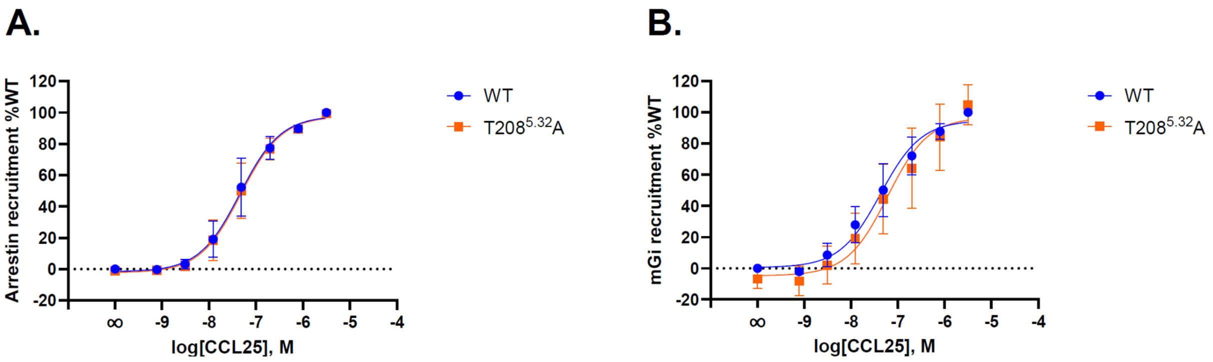


**Fig. S7. Previously reported G protein-biased ECL2 mutation in CCR9 is exactly WT-like in our experiments.** Concentration-dependent β-arrestin2 recruitment (**A**) and mGi recruitment (**B**) towards either CCR9 (blue) or CCR9 mutant $T208^{5.32}A$ (orange) following stimulation across a titration of CCL25 concentrations. Values represent mean ± SD of three independent experiments performed in duplicate, normalized to min and max WT response.

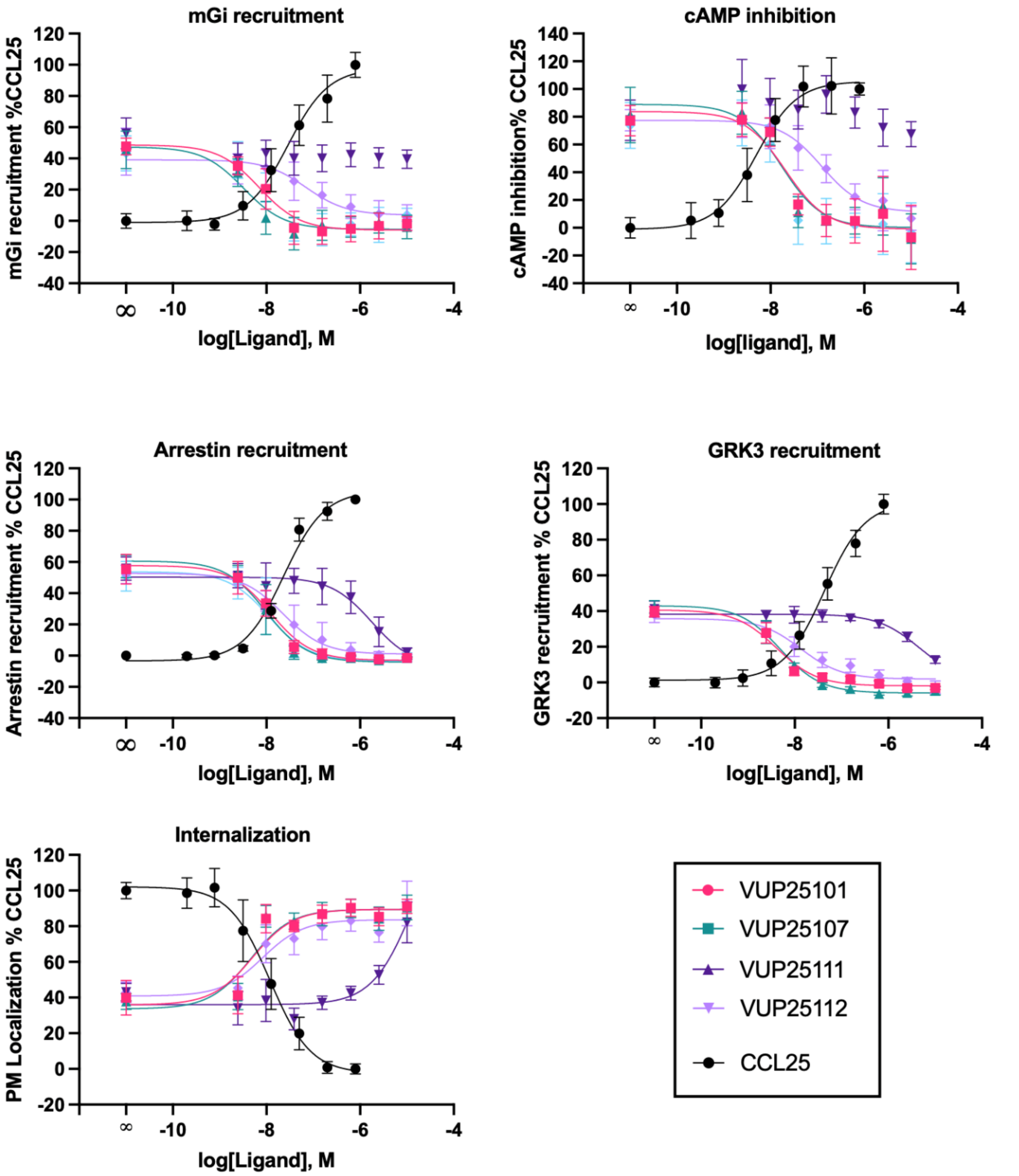


**Fig. S8. Overview of assays for comparison of the pharmacological properties of the miniproteins.** Results of the CCR9 pharmacological assays performed. Data presented is repeated from Figs. 4, 5, 6, and 7 and recompiled here for side-by-side comparison.

**Table S2. Statistics**

| Fig. | Condition: | N | Log EC/IC50 | | Statistical test |
|---|---|---|---|---|---|
| | | | P-value | Value ± SD | |
| 4B | **CCR9-β-Arrestin Recruitment:**<br>- VUP25101 as reference<br>- VUP25107<br>- VUP25111<br>- VUP25112 | 3 | <br>--<br>0.4870<br><0.0001<br>0.1059 | <br>-7.92±0.07<br>-8.01±0.24<br>-5.77±0.33<br>-7.56±0.41 | <br>Extra sum-of-squares F test |
| 4C | **ACKR4-β-Arrestin Recruitment:**<br>- VUP25101 as reference<br>- VUP25107<br>- VUP25111<br>- VUP25112 | 3 | <br>--<br>0.8320<br>0.0002<br><0.0001 | <br><br>-7.73 ±0.08<br>-7.71 ±0.04<br>N.A.<br>-6.84 ±0.37 | <br>Extra sum-of-squares F test |
| 5B | **CCR9 mGi Recruitment:**<br>- VUP25101 as reference<br>- VUP25107<br>- VUP25111<br>- VUP25112 | 3 | <br>--<br>0.0334<br>N.A.<br>0.0003 | <br>-8.17 ±0.21<br>-8.44 ±0.17<br>N.A.<br>-7.15 ±0.51 | <br>Extra sum-of-squares F test |
| 5D | **CCR9 cAMP inhibition:**<br>- VUP25101 as reference<br>- VUP25107<br>- VUP25111<br>- VUP25112 | 3 | <br>--<br>0.6073<br>0.0043<br>0.0008 | <br>-7.68 ±0.10<br>-7.75 ±0.25<br>N.A.<br>-6.98 ±0.25 | <br>Extra sum-of-squares F test |
| 6B | **CCR9-GRK3 Recruitment:**<br>- VUP25101 as reference<br>- VUP25107<br>- VUP25111<br>- VUP25112 | 3 | <br>--<br>0.9519<br><0.0001<br>0.1439 | <br>-8.40 ±0.04<br>-8.32 ±0.10<br>-5.33 ±0.36<br>-7.81 ±0.46 | <br>Extra sum-of-squares F test |
| 7B | **CCR9 Internalization:**<br>- VUP25101 as reference<br>- VUP25107<br>- VUP25111<br>- VUP25112 | 3 | <br>--<br>0.9526<br><0.0001<br>0.2647 | <br>-8.30 ±0.12<br>-8.35 ±0.16<br>-4.85 ±0.40<br>-8.04 ±0.33 | <br>Extra sum-of-squares F test |
| S1A | **CCR9-β-Arrestin Recruitment:**<br>- CCL25 as reference<br>- CCL25-ΔCT | 3 | <br>--<br><0.0001 | <br>-7.46 ±0.16<br>-7.90 ±0.20 | <br>Extra sum-of-squares F test |
| S1B | **ACKR4-β-Arrestin Recruitment:**<br>- CCL25 as reference<br>- CCL25-ΔCT | 3 | <br>--<br>0.6829 | <br>-7.85 ±0.25<br>-7.87 ±0.70 | <br>Extra sum-of-squares F test |
| S6C | **ACKR4 pIC50 CCL25/CCL20**<br>- CCL25<br>- CCL20 | | <br>--<br>0.0096 | <br>7.73 ± 0.08<br>6.06 ± 0.13 | <br>T-test |
| S7A | **CCR6:**<br>- VUP25101<br>- VUP25107<br>- VUP25111<br>- VUP25112 | 3 | <br>N.A. | <br>N.A.<br>-6.99 ±0.15<br>N.A.<br>-7.49 ±0.07 | <br>N.A. |

| | | | | | |
|---|---|---|---|---|---|
| **S7B** | **ACKR4:**<br>- VUP25101<br>- VUP25107<br>- VUP25111<br>- VUP25112 | 3 | <br>N.A. | <br>-6.06 ±0.13<br>-7.72 ±0.21<br>N.A.<br>-6.93 ±1.03 | N.A. |
